# Proteomic Analysis Reveals the Molecular Pathways Responsible for Solar UV-B Acclimation in High-altitude Malbec Berries

**DOI:** 10.1101/2023.10.20.562799

**Authors:** Leonardo A. Arias, Germán Murcia, Federico Berli, Flavio Muñoz, Ariel Fontana, Patricia Piccoli

## Abstract

Grapevine cultivation at high altitudes provides a viable option for producing premium quality wines in the context of climate change. This is primarily attributed to cooler temperatures, wider thermal amplitudes, and increased UV-B radiation. Although high UV-B levels can cause oxidative-stress, grape berries acclimate by generating UV-blocking anthocyanins and antioxidant compounds accumulated in the berry skins, thereby enhancing the organoleptic qualities and aging capacity of wine. This UV-B exclusion study examines how Malbec berries respond to solar UV-B at a high-altitude vineyard in Mendoza, Argentina (1350 m a.s.l.). The results showed that high solar UV-B acts both as a photomorphogenic signal and a stressor. The proteomic changes of berries exposed to +UV-B conditions indicate a decrease of photosynthesis and oxidative phosphorylation, coupled with an increase of glycolysis and tricarboxylic acid cycle as compensatory respiration pathways. Furthermore, numerous chaperones and proteins associated with the antioxidant system exhibited increased abundance to maintain cellular homeostasis. Lastly, veraison-stage berries exposed to +UV-B displayed an activation of the UVR8 signaling cascade and the phenylpropanoid pathway, resulting in higher concentration of phenolic compounds and more oxidation-resistant types of anthocyanins. This is the first report of field-grown grape berry proteomic modulation in response to solar UV-B, and it may have significant implications for the cultivation of high-quality wine grapes in both current and future climate scenarios.

**Significance:** 

## Introduction

The progression of climate change is prompting winegrape growers from regions with warmer growing seasons to explore more suitable environments for high-quality wine production, including higher altitude locations (Van Leeuwen et al., 2019, Arias et al., 2022). The unique environmental conditions found at higher altitudes, like increased solar radiation, cooler temperatures, and reduced atmospheric pressure, exert a significant influence on grapevine growth and development. This influence results in distinctive berry characteristics and wine quality (Berli et al., 2011; Martínez-Lüscher et al., 2017). Among these factors, solar ultraviolet-B (UV-B) radiation (280–315 nm) plays a crucial role in modulating grapevine physiology and biochemistry, particularly in high-altitude vineyards where UV-B levels are substantially higher than at sea level (Berli et al., 2010). According to its intensity and the interplay with other environmental factors, UV-B can act as a morphogenetic signal, regulating plant growth and development, or as a stressor, generating reactive oxygen species (ROS) and disrupting several cellular processes (Chen et al., 2022). Prior studies have demonstrated that solar UV-B radiation can trigger various molecular and biochemical changes in grapevine, including the accumulation of secondary metabolites like polyphenolic compounds and terpenoids (volatiles), which are essential for grapevine health, berry quality, and the organoleptic properties of wine (Berli et al., 2011, Gil et al., 2012).

Broadly, from an oenological perspective, grape berry polyphenolic compounds can be divided into anthocyanins and non-anthocyanins. Anthocyanins are responsible for wine color composition, contributing to the visual appeal of red wines. Non-anthocyanin polyphenolic compounds (NAPC) encompass a wide range of substances including flavonols, flavan-3-ols, stilbenes and phenolic acids. These compounds shape the overall flavor, astringency, and mouthfeel of wines (Berli et al., 2011).

UV-B radiation has been shown to modulate the expression of genes involved in the biosynthesis of these secondary metabolites, as well as those associated with the antioxidant system and stress responses (Berli et al., 2010; Martínez-Lüscher et al., 2017). These findings suggest the existence of a complex regulatory network underlying the grapevine response to UV-B. In a transcriptome study performed *in vitro*, Pontin et al. (2010) observed a UV-B-intensity dependent response in Malbec leaves. While high fluence rate UV-B exposure triggers general multi-stress defense response pathways, relatively low fluence rates (like those typically found in the field) promote UV-B protection. Furthermore, in an experiment simulating a climate change scenario, a higher UV-B regime was shown to mitigate some of the adverse effects caused by increased temperature and CO_2_, such as delaying premature ripening and reducing lipid peroxidation (Martínez-Lüscher et al., 2015).

It is worth pondering that a plant’s perception of solar UV-B integrates into a complex network of interactions downstream of multiple photoreceptors that sense a broad spectrum of light (Rai et al., 2021). However, most UV-B experimentation involves narrow-band UV-B supplementation on a white light background, and conducting long-term field experiments poses significant challenges due to the inherent variability and unpredictability of weather conditions that interplay, such as cloud cover (Palma et al., 2021; Rai et al., 2021). In recent years, authors have increasingly recognized the necessity to study physiologically relevant light conditions, or at least, improving the modeling of UV-B experimental conditions to mimic those found in vineyards (Rai et al., 2021). Supporting this perspective, experiments assessing the interplay between UV-B and different light spectra have shown that the quality of background light profoundly affects the capacity to both generate and scavenge ROS (Rácz and Hideg, 2021). Moreover, it influences the extent to which UV-B inhibits growth and the susceptibility to photoinhibition (Palma et al., 2021).

This study was aimed to elucidate the molecular pathways leading to biochemical changes in Malbec berry composition and in its physiological acclimation to high-altitude solar UV-B radiation. To achieve this, a UV-B exclusion experiment at a commercial high-altitude (1350 m a.s.l.) vineyard in Mendoza, Argentina was conducted. This region provides an invaluable opportunity to study very high solar UV-B regimes in relatively cloudless skies. The study employed label-free proteomics and STRING networks for data visualization, enabling valuable insights into the underlying molecular mechanisms governing the quality attributes of grape berries. By revealing the proteomic modulation in Malbec berries exposed to the intense solar UV-B radiation found at high altitudes, the study contributes to a deeper understanding of the complex interplay between environmental factors and the physiology of grapevine. This knowledge has the potential to impact viticultural practices and to enhance berry quality in the context of climate change challenges.

## Materials and Methods

### Plant Material and Experimental Design

The experiment was conducted during the 2020–2021 growing season using Malbec vines grown on their own roots in a high-altitude vineyard located in Gualtallary, Mendoza, Argentina (69°14′10″ W and 33°22′12″ S) at 1350 m a.s.l. Vine rows were oriented in a N-W to S-E direction, spaced 2,0 m between rows and 1.20 m between plants. Plants were pruned to 4 shoots and two clusters per shoot. Briefly, two distinct UV-B treatments were implemented: a low UV-B treatment (-UV-B) by using a 100 μm polyester cover capable of absorbing up to 90% of UV-B radiation at solar noon (Berli et al., 2008); while an uncovered UV-B treatment (+UV-B) was established as a control. For each treatment, a total of 21 plants were used. UV-B treatments started 20 days before veraison (stage 35, Coombe, 1995) and continued until harvest when berries reached 24 °Brix. Vines were maintained with no soil water restriction by drip irrigation.

### Light measurements

Photosynthetically active radiation (PAR) and UV-B irradiation were measured by a LI-250 light meter with a quantum sensor LI-190SA (Li-Cor Inc., Lincoln, NE, USA), and a PMA2200 radiometer with a PMA2102 detector (Solar Light Company Inc., Glenside, PA, USA) respectively.

Specifically, during the sunniest period of a mid-summer day, PAR (400-700 nm), UV-B (280-315 nm) were measured for +UV-B and -UV-B conditions from 10:00 am to 16:00 pm. The plastic covering effectively filtered out more than 80% of UV-B during this period, reaching an 89% reduction at noon (Fig. S1 a). Conversely, PAR was only reduced by up to 10% at solar noon (Fig. S1 b).

### Tissue collection

Sampling was carried out by selecting individual, sun-exposed berries from the clusters. For the pre-veraison stage, individual green berries were collected from clusters that reached 50% veraison. In contrast, for the veraison stage, firm red berries were harvested from clusters that had recently reached 100% veraison. We collected a minimum of 5 berries per condition and phenological stage from each of the 21 plants. Immediately after collection, samples were transported in coolers to the laboratory for weight and size measurements. The equatorial diameter of each berry was measured in ImageJ 1.53s (National Institutes of Health, USA, http://imagej.nih.gov/ij). Then, pulps were discarded by compressing the berries, and the skins were frozen using liquid nitrogen and kept at - 80 °C until further processing.

### Extraction of berry skin polyphenolic compounds

The extraction for each replicate was achieved by macerating 10 randomly selected berry skins per condition in a solution of 1% HCl-methanol at a 1:10 w/v ratio for 1 h at 70 °C, followed by three rounds of 5 min sonication. Subsequently, the solutions were centrifuged, and the supernatant collected for spectrometric measurements and chromatographic analysis.

### Standards, solvents and reagents

Standards of (+)-catechin (≥99%), (−)-epicatechin (≥95%), (+)-procyanidin B1 (≥90%), procyanidin B2 (≥90%), (−)-epigallocatechin (≥95%), (−)-gallocatechin gallate (≥98%), (−)-epicatechin gallate (≥95%), polydatin (≥95%), piceatannol (≥95%), (+)-ε-viniferin (≥95%), quercetin hydrate (95%), quercetin 3-β-d-galactoside (≥97%), quercetin 3-β-d-glucoside (≥90%), kaempferol-3-glucoside (≥99%), myricetin (≥96%), naringin (≥95%), 3-hydroxytyrosol (≥99.5%), caftaric acid (≥97%), ferulic acid (≥99%), gallic acid (99%), phlorizdin (≥99%) were purchased from Sigma–Aldrich (St Louis, MO, USA). Stock solutions were prepared in methanol at 1000 µg mL^−1^ concentration. Further dilutions were prepared in methanol and stored in dark-glass bottles at −20° C. For quantification of compounds by high performance liquid chromatography with diode array and fluorescence detection (HPLC-DAD-FLD), calibration standards were prepared in ultra-pure water (0.1% formic acid; FA)/Acetonitrile (MeCN) (95:5). HPLC-grade MeCN and FA were sourced from Mallinckrodt Baker Inc. (Pillispsburg, NJ, USA). Ultrapure water was procured from a Milli-Q system (Millipore, Billerica, MA, USA).

### Polyphenolic compounds profiling

Profiling of polyphenolic compounds was done by HPLC-DAD-FLD (Dionex Ultimate 3000 system, DionexSoftron GmbH, Thermo Fisher Scientific Inc., Germering, Germany). Anthocyanin determination was performed according to Urvieta et al. (2018), with minor adjustments. Briefly, a 500 μL aliquot of berry phenolic extract was dried by vacuum centrifugation and dissolved in 1 mL of initial mobile phase prior to chromatographic analysis. Anthocyanins were separated in a reversed-phase Kinetex C_18_ column (3.0× 100 mm, 2.6 μm) Phenomenex (Torrance, CA, USA). The mobile phase was composed of ultrapure H_2_O:FA:MeCN (87:10:3 v/v/v; eluent A) and ultrapure H_2_O:FA:MeCN (40:10:50 v/v/v; eluent B). Separation gradient was: 0 min, 10% B; 0-10 min, 25% B; 10-15 min, 31% B; 15-20 min, 40% B; 20-30 min, 50% B; 30-35 min, 100% B; 35-40 min, 10% B; 40-47 min, 10% B. Mobile phase flow, column temperature and injection volume were 1 mL min^−1^, 25 °C and 5 μL, respectively. Quantification was carried out by measuring peak area at 520 nm and the content of each anthocyanin was expressed as malvidin-3-glucoside equivalents, using an external standard calibration curve (1–250 mg L^−1^, r^2^ =0.997). The identity of detected anthocyanins was confirmed by comparison with the elution profile and identification of analytes performed in a previous research (Antoniolli et al., 2015). For non-anthocyanins compounds, phenolic extracts were analyzed according to analytical conditions reported by Ferreyra et al. (2021), using the same column as for anthocyanins. The mobile phases were an aqueous solution of 0.1% FA (solvent A) and MeCN (solvent B). The gradient was as follows: 0–1.7 min, 5% B; 1.7– 10 min, 30% B; 10–13.5 min, 95% B; 13.5–15 min, 95% B; 15–16 min, 5% B; 16–19, 5% B. The total flow rate was set at 0.8 mL min^-1^. The column temperature was 35 °C and the injection volume was 5 μL. The identification of non-anthocyanins was based on the comparison of the retention times of phenolic compounds in samples with those of authentic standards. External calibration was used as a quantification approach and linear ranges between 0.05 and 40 mg L^−1^ with coefficient of determination (r^2^) higher than 0.993 were obtained.

### Protein extraction and quantification

Protein fraction was extracted using the method previously described by Negri et al. (2008), with some modifications. Frozen berry skins (3 g) were finely powdered in liquid nitrogen using a pestle and mortar. The powder was then resuspended in 12 mL of extraction buffer [0.7 M sucrose (Cicarelli Labs, Santa Fe, Argentina), 0.5 M Tris-HCl pH 8 (MP Biomedicals, California, USA), 10 mM disodium EDTA salt (Promega, Madison, WI, USA), 1 mM PMSF (PhenylMethylSulfonyl Fluoride, Sigma-Aldrich Corp., St Louis, MO, USA), 0.2 % (v/v) β-mercaptoethanol (Sigma-Aldrich Corp., St Louis, MO, USA), protease inhibitor cocktail (Sigma-Aldrich Corp., St Louis, MO, USA) and PVPP (Sigma-Aldrich Corp., St Louis, MO, USA)] and incubated at 4°C under shaking for 10 min. Proteins were extracted by the addition of an equal volume of ice-cold Tris-buffered phenol pH 8 (Sigma-Aldrich Corp., St Louis, MO, USA). The sample was shaken for 30 min at 4°C, incubated for 2 h at 4°C and finally centrifuged at 5000 g for 20 min at 4 °C to separate the phases. Then, 10 mL of the upper phenol phase was collected, and the proteins were precipitated by the addition of 40 mL of ice-cold 0.1 M ammonium acetate in methanol.

The sample was vortexed briefly and finally maintained at −20 °C overnight. Precipitated proteins were recovered by centrifuging at 13000 g for 30 min at 4 °C, then washed again with cold methanolic ammonium acetate and three additional times with cold 80% (v/v) acetone (Sintorgan, Bs As, Argentina). The final pellet was dried at room temperature and resuspended in 500 µL of [7 M urea (Promega, Madison, WI, USA), 2 M thiourea (Sigma-Aldrich Corp., St Louis, MO, USA), 4 % (v/v) IGEPAL (Sigma-Aldrich Corp., St Louis, MO, USA) and 50 mg mL^-1^ DTT (Sigma-Aldrich Corp., St Louis, MO, USA)] buffer. Finally, the sample was centrifuged at 13000 g for 3 min and the supernatant stored at −80 °C until further use. The protein concentration was determined by the Bradford assay (Bio-Rad Laboratories Inc, California, USA).

### nano-HPLC-mass spectrometry protein analysis

Protein samples were boiled (95 °C, 5 min) in Laemmli buffer and 50 µg of total protein was run in 10 % SDS-PAGE. Once the proteins ran 2 cm into the separating gel, the running was stopped. The gel was stained according to colloidal Coomassie Brilliant Blue G-250 (cCBB) procedure (Neuhoff et al., 1988), and the dried fragments of the gel corresponding to each treatment replica were sent to the Proteomics Core Facility CEQUIBIEM, Buenos Aires, Argentina. Proteins were reduced with 10 mM DTT for 45 min at 56 °C and alkylated with 50 mM iodoacetamide for 45 min in darkness. Proteins were digested overnight with sequencing-grade modified trypsin (Promega). Then, the samples were lyophilized with SpeedVac and resuspended with 30 µL of 0.1 % trifluoroacetic acid. Zip-Tip C_18_ (Merck Millipore) columns were used for desalting. Resulted peptides were separated in a nano-HPLC (EASY-nLC 1000, Thermo Fisher Scientific, Germany) coupled to a mass spectrometer with Orbitrap technology (Q-Exactive with High Collision Dissociation cell and Orbitrap analyzer, Thermo Fisher Scientific, Germany). Peptides were ionized by electrospray (EASY-SPRAY, ThermoScientific), at a voltage of 1.5 to 3.5 kV.

### Proteomics data analysis

Proteome Discoverer 2.2 software (ThermoScientific, Germany) was used to match the identity of peptides to the grapevine reference proteome set from uniprot (Vitis vinifera-UP000009183-Uniprot). The raw intensity values obtained from the mass spectrometry data were normalized using the total ion current normalization method. The critical search parameters were as follows: precursor ion mass tolerance of 10 ppm, fragment mass tolerance of 0.05 Da, trypsin enzyme with a maximum of two missed cleavages allowed, variable modifications including oxidation and carbamidomethylation of cysteine, and a minimum of two peptides identified per protein. Missing values in the dataset were calculated by the mean imputation method. Only proteins with high confidence and a percolator q-value lower than 0.01 were considered for further analyses. To identify differentially abundant proteins, a two-sample t-test was performed with a significance threshold of p < 0.05. The fold changes of protein abundances were log_2_ transformed for further analysis.

Volcano plots were created using Perseus 1.6.6 software platform (Max-Planck-Institute of Biochemistry, Germany). The heatmaps used for UV-B treatments comparison were designed using pheatmap package (Kolde, 2019) in R 4.1.1 (R Core Team, 2023). Bubble plots showing GO terms were designed using the ggplot2 package (Wickham et al., 2022) in R 4.1.1 and custom R scripts. Selected GO terms for biological processes were retrieved from STRING functional enrichment of stringApp 1.7 (Doncheva et al., 2019) in the Cytoscape 3.9.1 open software platform (Shannonet al., 2003) with the grapevine reference genome as background and avoiding redundancy.

Full STRING networks were constructed with stringApp for Cytoscape, considering only significantly regulated proteins without a fold change cutoff. Instead, log_2_ of the abundance fold change for each protein was color coded as a heatmap inside each node. Network clustering was performed with the MCL algorithm in stringApp for Cytoscape, with an inflation value of 4. Each cluster was labeled with their most representative GO term for biological function, which was retrieved by setting the redundancy cutoff to 0 in STRING functional enrichment.

## Results and Discussion

This work constitutes the first label-free proteomic study of the UV-B response in field-grown grapevine<u>s</u>. The effects of high-altitude solar UV-B radiation on the proteomic and polyphenolic profile of Malbec berry skin berries were assessed at two ripening stages: pre-veraison and veraison. All comparisons were made to highlight the uncovered, high UV-B (+UV-B) condition as compared to the covered, low UV-B (-UV-B), to better assess the contribution of solar UV-B to the modulated pathways.

### High-altitude UV-B significantly increases the accumulation of phenolic compounds

A total of 21 non-anthocyanin phenolic compounds (NAPCs) were identified by HPLC-DAD-FLD, distributed among procyanidins, prodelphinidins, stilbenes, flavonols, hydroxycinnamic acids, hydroxybenzoic acids and dihydrochalcones. UV-B exposure markedly increased the concentration of all the NAPCs in pre-veraison berries, ranging from 200 to 500% (Table 1), confirming that UV-B is a major contributor to the accumulation of NAPCs in pre-veraison. This is also supported by spectrometric measurements showing a significant increase in absorbances corresponding to total polyphenols (280 nm) by about 67% and total UV-absorbing compounds (305 nm) by 35% for the uncovered +UV-B condition as compared to −UV-B (Fig. S2). These findings are consistent with the findings of Berli et al. (2011) and Marfil et al. (2019), who reported that UV-B radiation increased the biosynthesis of NAPCs such as flavonols, hydroxycinnamic acids, and stilbenes. During veraison, the onset of ripening characterized by the appearance of color in varieties for red wine, the concentration of procyanidins and prodelphinidins in berry skins sharply decreased regardless of the UV-B exposure regime, although in the uncovered +UV-B grapes it decreased further (Table 1). This phenomenon has been described previously as a result of the dynamic between biosynthesis, polymerization and degradation that occurs in berry skins during ripening, as these compounds are the precursors of tannins (Downey et al., 2003). These authors found that compounds such as catechin, epicatechin and epicatechin-gallate have a peak of concentration between fruit set and two weeks before veraison, decreasing rapidly during veraison. Moreover, this dynamic has been found to be cultivar dependent and influenced by abiotic factors, including UV-B, as thoroughly reviewed by Blancquaert et al. (2019). A significantly higher UV-B regime could prematurely cause a net decrease in these NAPCs during veraison by promoting polymerization into other compounds, forming associations with cellular components like the cell wall, shifting the biosynthetic pathway into anthocyanin production, being oxidized by UV-B, or a combination of those factors (Blancquaert et al., 2019).

**Table 1.** Non-anthocyanin polyphenolic compounds found in berry skins. Concentration expressed as ng of compound per g of fresh berry skin. ND: not detected. Two-way ANOVA with Tukey contrasts was performed (DFn=1, DFd=8). Letters represent significantly different values, with p<0.05

| Compound | Pre-veraison -UV |  | Pre-veraison +UV |  | Veraison -UV |  | Veraison +UV |  |
| --- | --- | --- | --- | --- | --- | --- | --- | --- |
| Procyanidins |  |  |  |  |  |  |  |  |
| (+)-catechin | 192.8 ±54.9 | a | 661.1 ±60.3 | b | 24.1 ±4.4 | c | 17.9 ±3.6 | c |
| (-)-epicatechin | 38.3 ±4.9 | a | 82.2 ±1.8 | b | 14.5 ±0.3 | c | 8.1 ±0.1 | d |
| Procyanidin B1 | 1266.6 ±133.6 | a | 8929.5 ±498.4 | b | 89.3 ±6.7 | c | 68.7 ±1.5 | d |
| Procyanidin B2 | 249.2 ±87.9 | a | 586 ±55.6 | b | 59.2 ±2.3 | c | 29.4 ±1.3 | d |
| (-)-epigallocatechin | 223.2 ±85.9 | a | 642.7 ±108.7 | b | 1303.3 ±113.6 | c | 3296.5 ±1106.6 | c |
| Prodelphinidins |  |  |  |  |  |  |  |  |
| (-)-gallocatechin gallate | 273.1 ±35.5 | a | 1013.9 ±80.7 | b | 168.2 ±13.2 | c | 76.3 ±8.7 | d |
| (-)-epicatechin gallate | 184.8 ±41.4 | a | 444.8 ±59.8 | b | 98 ±4.8 | c | 48.6 ±4.6 | d |
| Stilbenes |  |  |  |  |  |  |  |  |
| Polydatin | 90.4 ±9.2 | a | 169.1 ±6.9 | b | 36.2 ±7.9 | c | 7.9 ±2.1 | d |
| Piceatanol | 9.9 ±1 | a | 25.9 ±0.8 | b | 18.8 ±0.5 | c | 9 ±0.1 | a |
| E-viniferin | 10 ±1.6 | a | 29.1 ±5 | b | 12.3 ±1.1 | a | 6.2 ±0.5 | c |
| Flavonoles |  |  |  |  |  |  |  |  |
| Quercetin | 156.7 ±14.2 | a | 410.1 ±47.6 | b | 179.4 ±19.2 | c | 214.9 ±25 | c |
| Quercetin-3-galactoside | 207 ±64.7 | a | 650 ±47.1 | b | 79.1 ±8.1 | c | 77.8 ±2.3 | c |
| Quercetin-3-glucoside | 7.2 ±2.3 | a | 25.6 ±0 | b | 4.7 ±0.7 | a | 1.7 ±0.6 | c |
| Kaempferol-3-glucoside | 130.6 ±27.2 | a | 333.5 ±19.2 | b | 136.6 ±10.1 | a | 173.5 ±13 | a |
| Myricetin | 368.3 ±175.1 | a | 1389.6 ±52.2 | b | 264.7 ±22.3 | a | 268.8 ±15.1 | a |
| Naringin | 17.2 ±4.6 | a | 53.1 ±1 | b | ND | c | ND | c |
| Hydroxycinnamic acids and Hydroxytyrosol |  |  |  |  |  |  |  |  |
| OH-Tyrosol | 196.7 ±27.4 | a | 439.7 ±44.5 | b | 175.5 ±16.4 | a | 177.6 ±15 | a |
| Caftaric acid | 89.5 ±21.2 | a | 306.8 ±12.6 | b | 53.4 ±2.5 | a | 33 ±5 | c |
| Ferulic acid | 18.7 ±7.5 | a | 45.3 ±6.5 | b | ND | c | 2.6 ±0.7 | d |
| Hydroxybenzoic acid |  |  |  |  |  |  |  |  |
| Gallic acid | 24.9 ±9.4 | a | 56.3 ±3.4 | b | ND | c | ND | c |
| Dihydrochalcone |  |  |  |  |  |  |  |  |
| Phloridzin | 4.2 ±6 | a | 20.8 ±2.6 | b | 0.8 ±1.2 | a | ND | a |

Anthocyanins accumulated significantly more in the +UV-B condition during veraison, ranging from 20 to 48% increase as compared to −UV-B (Table 2). Total anthocyanins were also spectrophotometrically measured by absorbance at 546 nm, showing an increase of about 35% for the +UV-B condition over the −UV-B (Fig. S2). This is a very well reported response of grape skins to UV-B (Berli and Bottini, 2013). No anthocyanins were detected by spectrophotometry or HPLC-DAD-FLD in the pre-veraison grapes, as we purposely selected completely green grapes that had not yet undergone veraison.

**Table 2.**
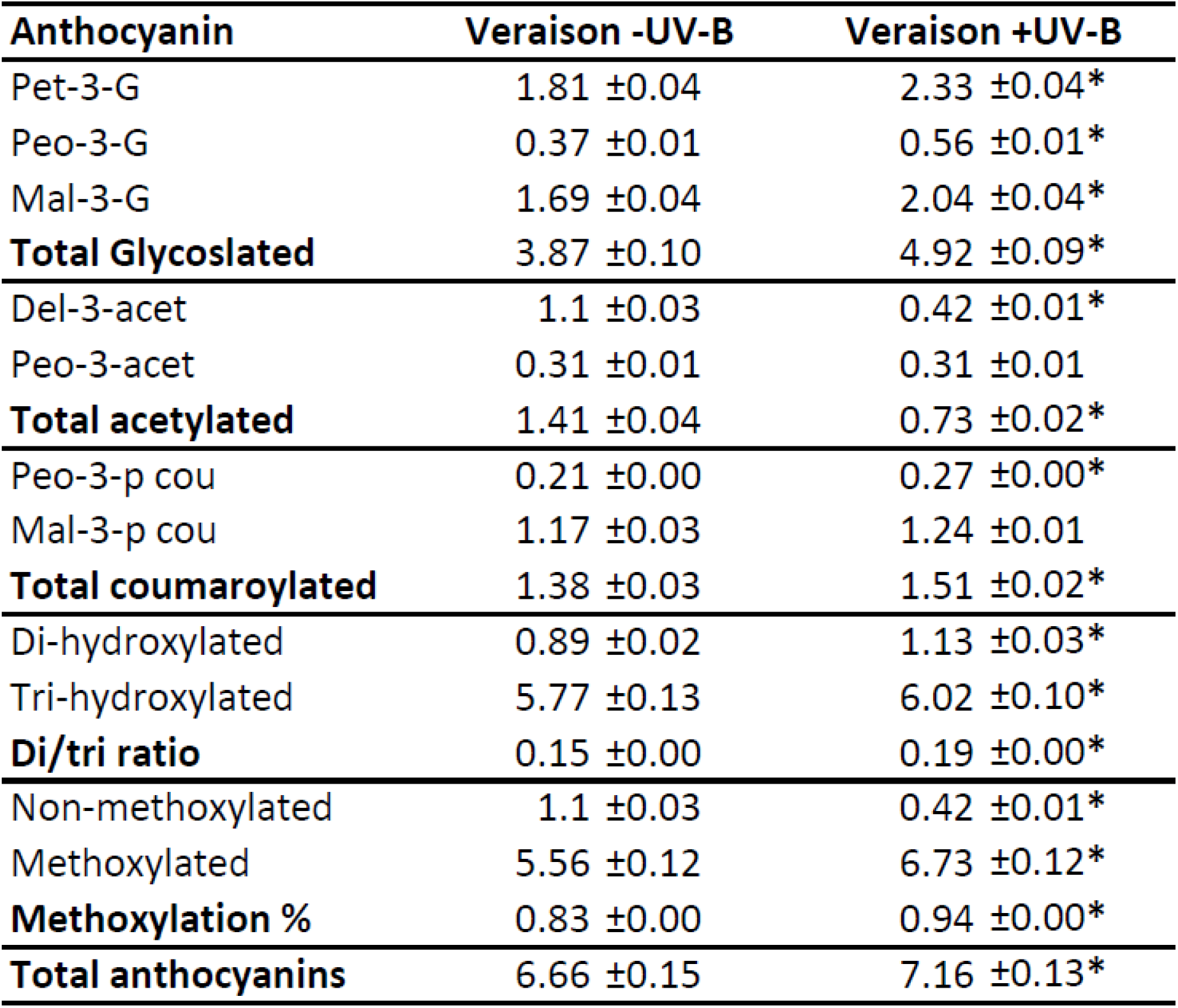
Anthocyanin compounds found in berry skins. Concentration expressed as μg of anthocyanin per g of fresh berry skin. One-way ANOVA with Tukey contrasts was performed. Asterisks represent significantly different values, with p>0.05 in a +UV-B vs –UV-B comparison.

| **Anthocyanin** | **Veraison -UV-B** | **Veraison +UV-B** |
| --- | --- | --- |
| Pet-3-G | 1.81 ±0.04 | 2.33 ±0.04* |
| Peo-3-G | 0.37 ±0.01 | 0.56 ±0.01* |
| Mal-3-G | 1.69 ±0.04 | 2.04 ±0.04* |
| **Total Glycoslated** | 3.87 ±0.10 | 4.92 ±0.09* |
| Del-3-acet | 1.1 ±0.03 | 0.42 ±0.01* |
| Peo-3-acet | 0.31 ±0.01 | 0.31 ±0.01 |
| **Total acetylated** | 1.41 ±0.04 | 0.73 ±0.02* |
| Peo-3-p cou | 0.21 ±0.00 | 0.27 ±0.00* |
| Mal-3-p cou | 1.17 ±0.03 | 1.24 ±0.01 |
| **Total coumaroylated** | 1.38 ±0.03 | 1.51 ±0.02* |
| Di-hydroxylated | 0.89 ±0.02 | 1.13 ±0.03* |
| Tri-hydroxylated | 5.77 ±0.13 | 6.02 ±0.10* |
| **Di/tri ratio** | 0.15 ±0.00 | 0.19 ±0.00* |
| Non-methoxylated | 1.1 ±0.03 | 0.42 ±0.01* |
| Methoxylated | 5.56 ±0.12 | 6.73 ±0.12* |
| **Methoxylation %** | 0.83 ±0.00 | 0.94 ±0.00* |
| **Total anthocyanins** | 6.66 ±0.15 | 7.16 ±0.13* |

Grape berry anthocyanins branch into two groups according to their level of hydroxylation, given by the activity of either flavonoid-3-hydroxylase (F3H), generating the di-hydroxylated cyanidin derivatives, or flavonoid 3’5’ hydroxylase (F3’5’H), generating the tri-hydroxylated delphinidin derivatives (Castellarin et al., 2007). These can be further methoxylated by the activity of o-methyltransferase (OMT), yielding peonidin from cyanidin, and petunidin or malvidin from delphinidin (Castellarin et al., 2007). The di/tri-hydroxylation ratio at maturity is reported to increase as a response to UV-B, favoring the more antioxidant di-hydroxylated forms (Berli et al., 2011) due to a preferential activation of F3’H over F3’5’H genes (Martínez-Lüscher et al., 2014).

In these results, +UV-B condition indeed increased the concentration of the di-hydroxylated peonidins by about 20%, resulting in a 27% increase of the di/tri ratio (Table 2), which is in line with previous reports of Berli et al. (2011). Moreover, +UV-B increased the methoxylated forms of anthocyanins (petunidin, peonidin and malvidin) from 83%, typical for red varieties grown at lower altitudes (Castellarin et al., 2007), to 94%.

### Malbec grape skin proteome is modulated by UV-B

After filtering by homology and statistical confidence, 1687 proteins were identified in at least two out of three replicates for pre-veraison and 1988 for veraison (Fig. 1). A post-hoc one-way ANOVA for pre-veraison and veraison data sets revealed 189 and 246 proteins respectively with significantly modified abundances caused by the UV-B exclusion treatment (Table S1), all of which were considered for further analyses. Volcano plots were constructed for each phenological stage, showing the differentially expressed proteins over the 5% confidence threshold line (Fig. 1). Heatmaps made through hierarchical clustering analysis show that protein abundances display significantly distinct patterns for both phenological stages and UV-B conditions (Fig. 2).

**Figure 1.**
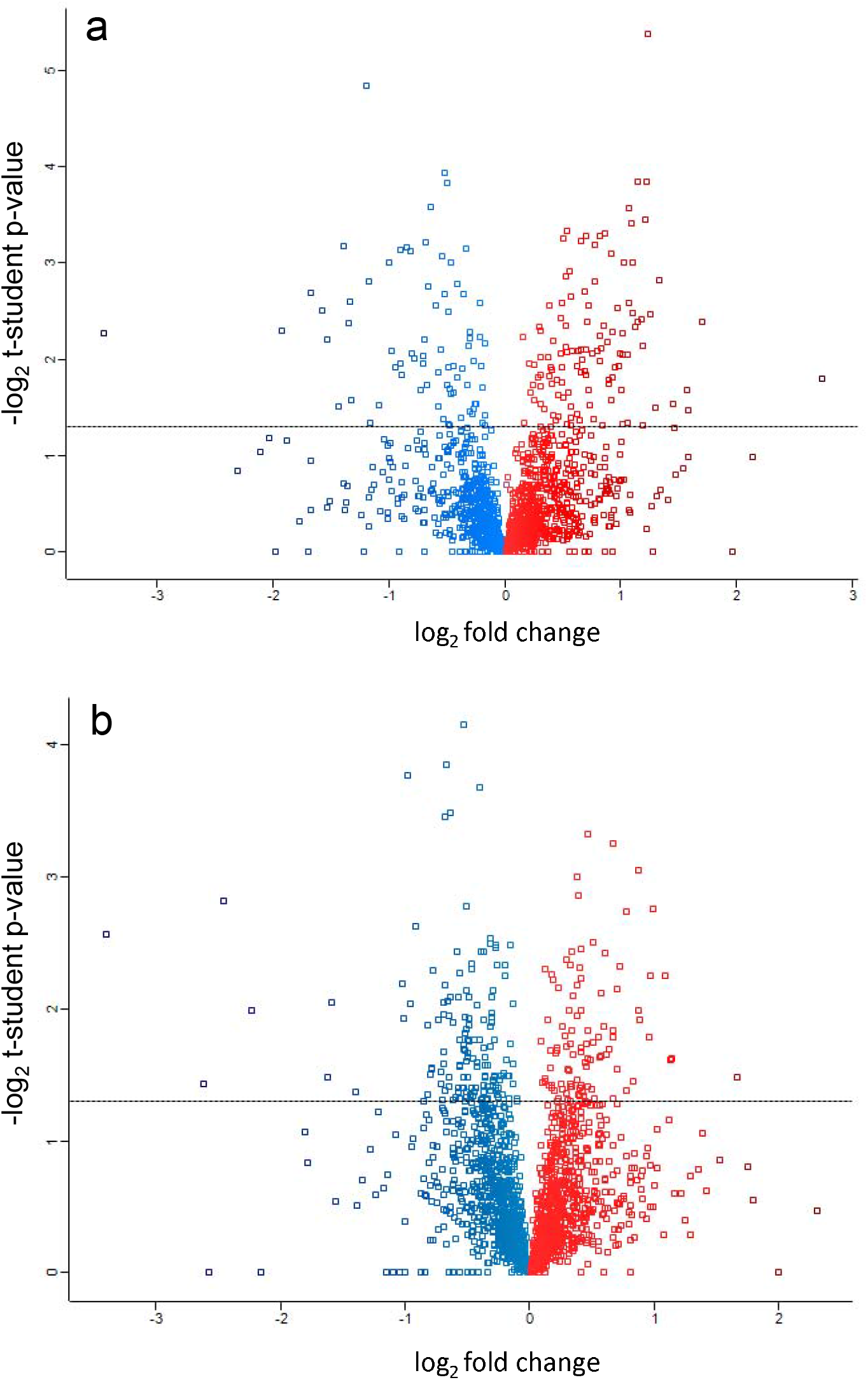
Volcano plots displaying the observed genome of grape skins subjected to UV-B exclusion during pre-veraison (a) and veraison (b). Dotted line marks the log_2_ transformed p=0.05 confidence threshold. Blue: reduced abundance; Red: increased abundance.

**Figure 2.**
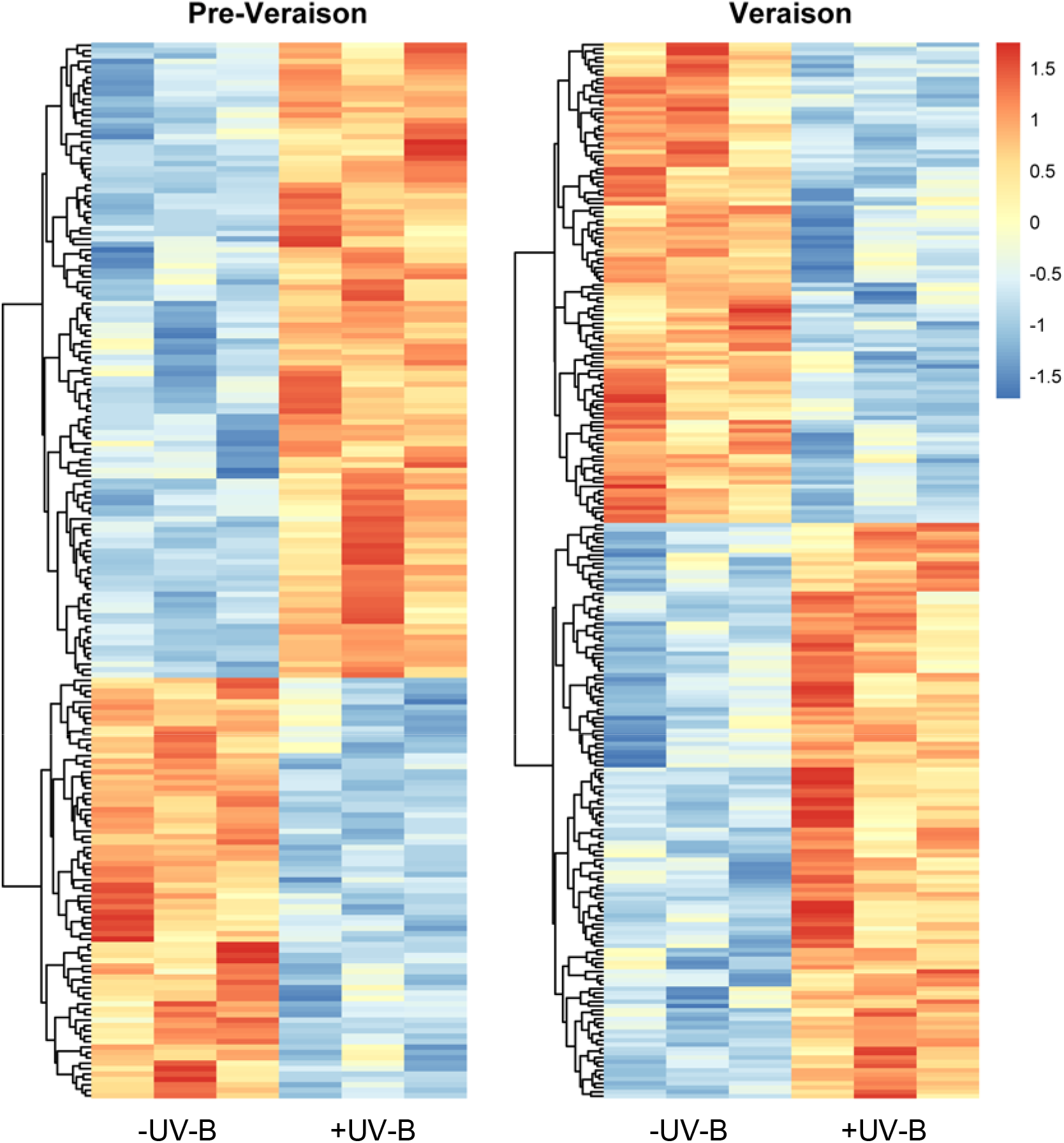
Heatmaps displaying hierarchical clustering of the three replicates corresponding to all differentially abundant proteins affected by the UV-B exclusion treatment. Blue: reduced abundance; Red: increased abundance.

The difference in the number of both detected and regulated proteins in pre-veraison and veraison could be a consequence of the unique set of biological processes occurring during these two phenological stages. In an RNA-Seq analysis of four developmental stages of Shiraz grape, Sweetman et al. (2012) showed how the transcriptome pattern is highly dependent on phenological stage. This is also supported by the distinct cluster association shown in the heatmaps for each stage (Fig. 2).

### Gene ontology

A gene ontology (GO) analysis was performed on the significantly regulated proteins for both phenological stages. The complete list of GO terms for biological processes (Table S2) below the 5% FDR (false discovery rate) was manually parsed for significance, enrichment strength (log_10_ (observed / expected)) and avoiding redundancy. Then a bubble plot was constructed with the most relevant terms (Fig. 3).

**Figure 3.**
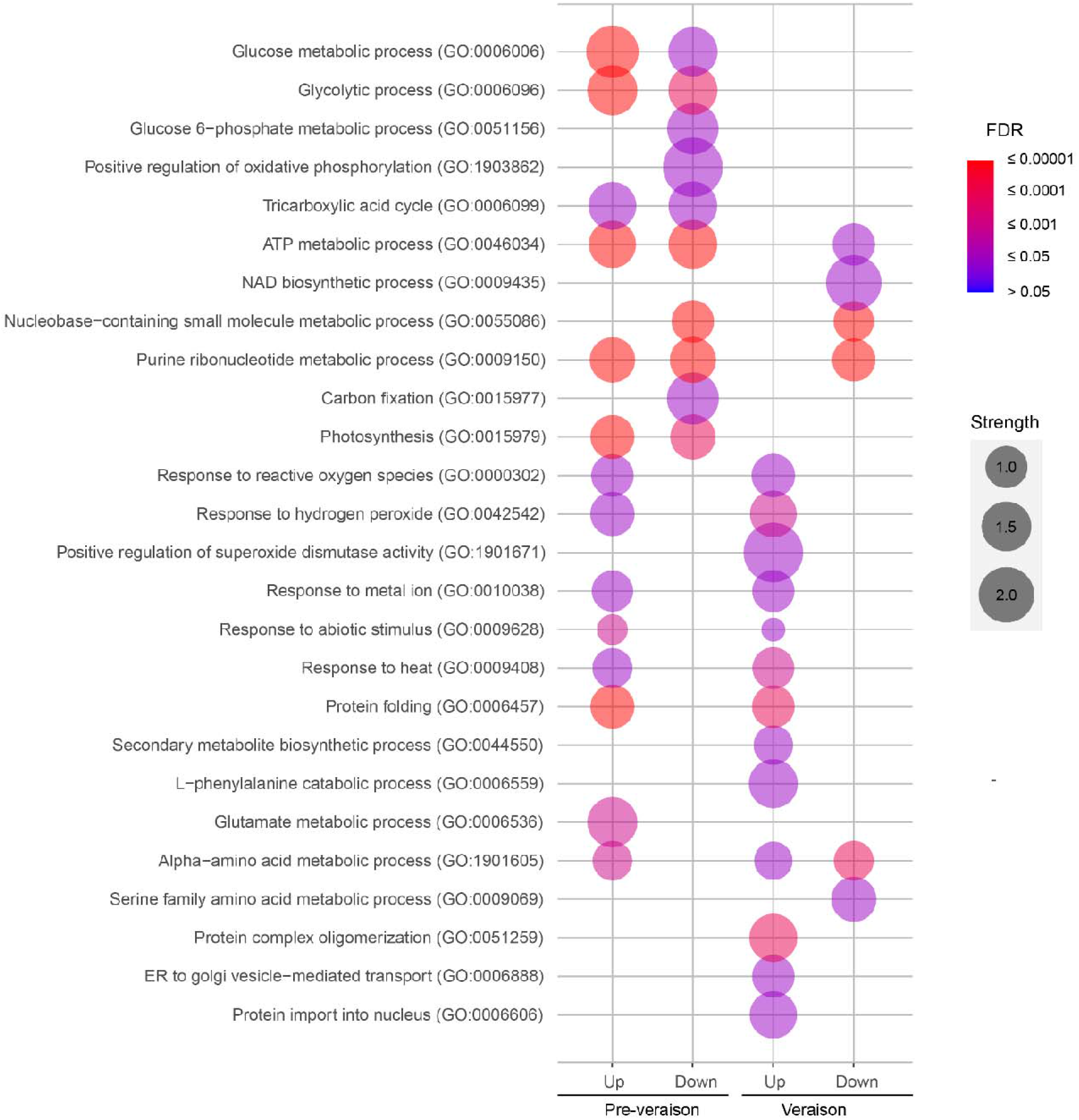
Bubble plot representing differentially abundant proteins into their Biological Process GO terms. Strength (Log10 (observed / expected) is a measure of functional enrichment.

The bubble plot highlights similarities and differences in the solar UV-B response in pre-veraison and veraison in terms of biological processes. Berries of both stages strongly favored abiotic stress categories, such response to hydrogen peroxide (GO:0042542), response to metal ion (GO:0010038), response to heat (GO:0009408) and protein folding (GO:0006457) (Fig. 3). This is an indication that the UV-B dose received by the uncovered berries was not only photomorphogenic but acted as a stressor. Although this is not often the case for ambient UV-B levels, as reviewed by Hideg et al. (2013), a high enough dose of UV-B will cause oxidative damage. This happens whenever plants are supplemented with high-fluence UV-B, as reviewed by Frohnmeyer and Staiger (2003). The same has been demonstrated for high-fluence UV-B supplemented Malbec leaves (Pontin et al., 2010), and for very high-altitude solar UV-B in Malbec vines (Berli et al., 2012).

Primary metabolism was also regulated in both stages, but in different ways. In pre-veraison, glycolytic process (GO:0006094), tricarboxylic acid cycle (TCA, GO:0006099), ATP metabolic process (GO:0046034) and photosynthesis (GO:0015979) categories, included proteins with both increased and reduced abundance, while glucose-6-phosphate metabolic process (GO:0051156), positive regulation of oxidative phosphorylation (GO:1903862) and carbon fixation (GO:0015977) included only proteins with reduced abundance (Fig. 3). Although UV-B as a photomorphogenic signal does not necessarily alter primary metabolism, when UV-B intensity is strong enough to be a stressor, cells can undergo pathway reprogramming to maintain their homeostasis (Kusano et al., 2011, Hideg et al., 2013). To cope with this stress, plants may up-regulate genes involved in glycolysis and the TCA cycle to increase ATP production and provide the necessary energy for the activation of various defense mechanisms, such as antioxidant enzyme systems (Frohnmeyer and Staiger, 2003). Note that pre-veraison berries were harvested still green, hence being able to carry out carbon fixation and photosynthesis. These processes have been shown to be impaired by UV-B in plants in general (Kataria et al., 2014), and in grapevine (Berli et al., 2012), by the combined effects of direct photosystem inhibition and impaired net gas exchange rate from UV-B-triggered lower stomatal conductance.

In veraison, on the other hand, there was a reduced abundance of proteins from ATP metabolism (GO:0046064) and NAD biosynthesis (GO:0009435) pathways, and a simultaneous increased abundance of the secondary metabolite biosynthesis (GO:0044550) and L-phenylalanine catabolic process (GO:0006559), which equates to the phenylpropanoid pathway (Fig. 3). This corresponds to the shift towards secondary metabolism that occurs at this stage, which is enhanced by UV-B as described by Martínez-Lüscher et al. (2016). Interestingly, authors pointed out that this could be desirable for wine production in warmer climates, as it prevents precocious, heat-related sugar maturity at harvest. Lastly, two categories related to retrograde intracellular transport (GO:0006888) and nucleocytoplasmic transport (GO:0006606) presented proteins with increased abundance only in veraison (Fig. 3), likely aiding the translocation of transcription factors and other messenger molecules during UV-B signaling.

### STRING networks

The UniProt codes of the 189 and 246 differentially abundant proteins for pre-veraison and veraison respectively were introduced into stringApp for Cytoscape, creating two full networks of protein interaction (Fig. S3 and Fig. S4). Log_2_ converted fold-change for each protein was then color coded as a heatmap inside each node. Pre-veraison and veraison networks displayed 192 nodes and 1819 edges, and 246 nodes and 789 edges, respectively. Both full networks were then clustered with the MCL algorithm inside stringApp, and the main GO term for biological process was used as a label for each cluster.

The STRING network analysis conducted in this study revealed the prominent clusters associated with the UV-B response during both the pre-veraison and veraison stages. The integration of STRING network analysis on top of gene ontology enabled the identification of both individual proteins and hidden molecular interaction networks involved in the complex response of Malbec grapes to high-altitude solar UV-B.

### Pre-Veraison STRING clusters

Pre-veraison STRING network separated into eight clusters with at least four nodes each, related to primary metabolism, peptide metabolism, protein folding, redox homeostasis and microtubule-based processes (Fig. 4).

**Figure 4.**
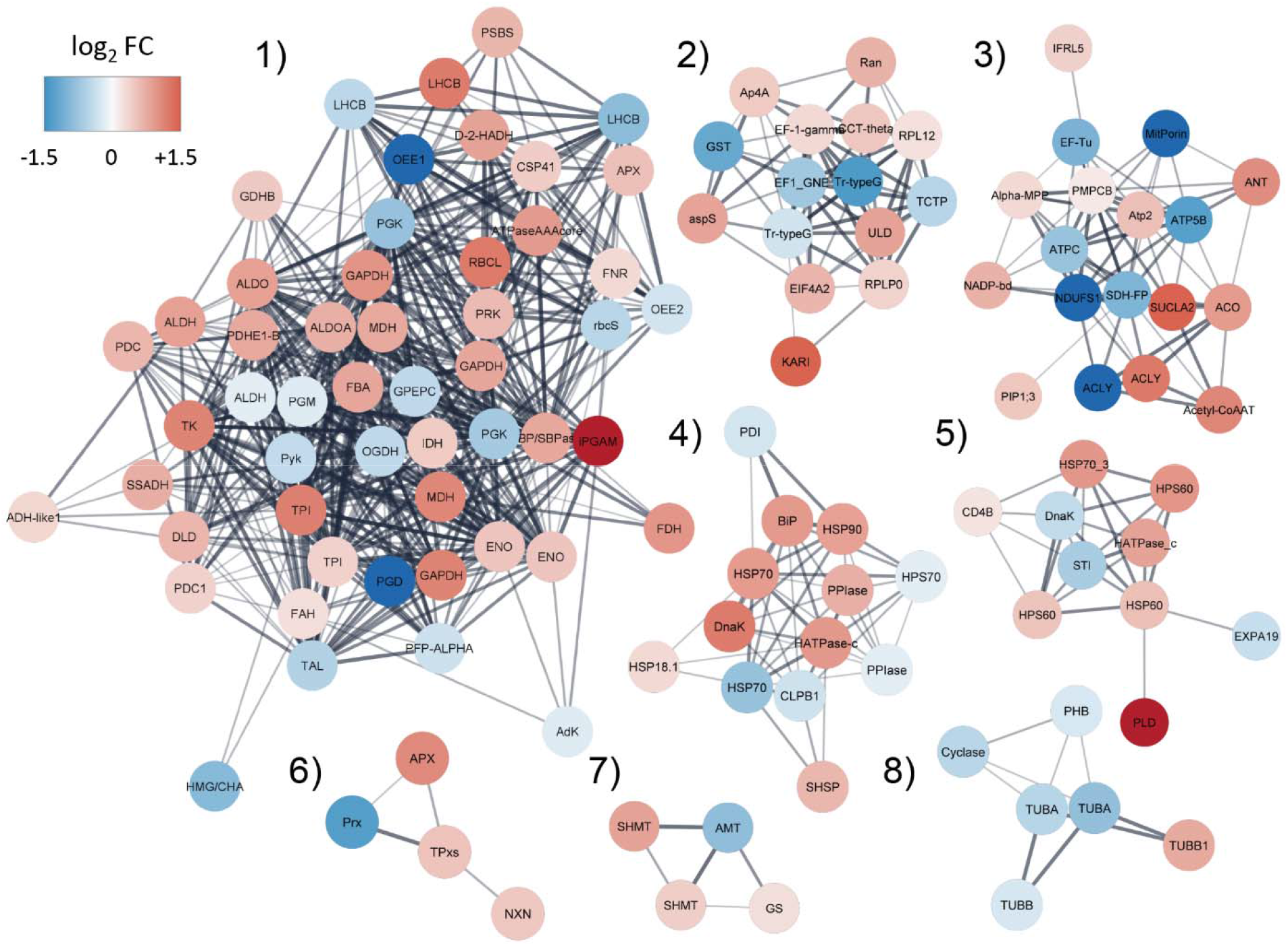
Principal STRING interaction clusters of significantly regulated proteins in pre-veraison grape skins. The full string network was clustered using the MCL app in Cytoscape with an inflation value of 4. All clusters with at least 4 nodes are shown, colored with their log_2_ of abundance fold change as a heatmap. Representative GO terms for biological process were retrieved using STRING enrichment for each cluster with a redundancy cutoff of 0.

1. Generation of precursor metabolites and energy
2. Peptide metabolic process
3. ATP metabolic process
4. Protein folding
5. Protein folding
6. Cell redox homeostasis
7. Glycine metabolic process
8. Microtubule based process

### Primary metabolism

Clusters 1 and 3, with a total of 68 proteins species involved in primary metabolism, indicate that this was the most regulated process among those affected by high UV-B in pre-veraison. Cluster 1 corresponded with the GO term “generation of precursor metabolites and energy” (GO:0006091), while cluster 3 associated with the GO term “ATP metabolic process” (GO:004e6034). The increased abundance of one or more species of triosephosphate isomerase (TPI, D7TLU7 and F6I134), glyceraldehyde-3-phosphate dehydrogenase (GAPDH, D7UDC9, F6H409 and F6GSG7), enolase (ENO, D7T227 and F6HKH3), pyruvate decarboxylase (PDC, D7TJI9 and F6GY71), pyruvate dehydrogenase (PDHE1, F6I1P0), aconitase (ACO, D7T7Y3), succinate-CoA ligase (SUCLA2, A5BF93), ATP citrate lyase (ACLY, D7SYK8), acetyl-CoA acetyltransferase (Acetyl-CoAAT, F6HHQ7), and malate dehydrogenase (MDH, F6GSG7 and D7FBC0), which are key enzymes in glycolysis and the tricarboxylic acid cycle (TCA) (Plaxton, 1996), suggests an increased energy demand under high UV-B conditions.

Additionally, cluster 1 exhibited the decreased abundance of photosynthetic proteins such as oxygen-evolving enhancer proteins (OEE1, F6I229 and OEE2, A5B1D3), as well as two species of light-harvesting chlorophyll protein complex (LHCB, F6GVX0 and F6HKS7). This corresponds with a general inactivation of photosynthesis that plants undergo as a response to high UV-B. As reviewed by Kataria et al. (2014), a high dose of UV-B can directly inactivate LHCB and OEE subunits, ultimately causing the inhibition of photosystem II. On the other hand, photosystem II subunit S (PSBS), known to have an important role in light acclimation and cross-tolerance to UV, displayed increased abundance (Górecka et al., 2020).

Lastly, cluster 3 displayed the reduced abundance of two ATP synthase subunits (ATPB, F6GTT2 and ATPC, F6H7M1) and the complex II subunit succinate dehydrogenase flavoprotein (SDH-FP, D7SPF1), indicating an impairment in the electron transport chain (ETC) process of ATP generation.

ETC is a major source of ROS in plants, primarily produced at complex I and III when the accumulated reduced intermediates transfer their electrons to molecular oxygen, generating superoxide and singlet oxygen (Jacoby et al., 2018). Typically, when ETC is blocked during stress, electrons can be transferred through several alternative points, such as alternative NAD(P)H-dehydrogenases and alternative oxidase (AOX) in order to prevent excess ROS and recycle the electron carriers (Van Aken et al., 2009; Schertl and Braun, 2014). For example, the down-regulation of the δ-subunit reduces ATP synthase abundance, impairing respiration, while simultaneously inducing glycolysis and non-energy conserving respiration through AOX (Geisler et al., 2012). In this study, although no AOX was detected in the observed proteome, a putative alternative NAD(P)H-dehydrogenase (F6HL96) with increased abundance was identified.

It is worth noting that this work focused on extracting soluble proteins, and consequently ETC proteins might be differentially represented in our extracts according to their tendency to remain associated with membranes. A proteomic study focusing on lipid soluble proteins would likely provide more reliable data towards unveiling ETC regulation during UV-B stress.

### Peptide metabolism

Cluster 2, identified with the GO term “peptide metabolic process” (GO:0006518) included proteins responsible for the fine-tuning of peptide biosynthesis during stress. The abundance of two ribosomal proteins (RPL12 and RPLP0), an elongation factor (EF-1), a eukaryotic initiation factor (EiF4A2), a chaperone (CCT-theta), a Ran-GTPase related to protein transport, and an aspartyl-tRNA synthetase (aspS) was increased. Ribosomal proteins not only can be tissue-specific and developmentally regulated, but they can often respond to environmental stimuli to specifically direct translation towards pertinent processes (Ferreyra et al., 2010). This is the case of RPLP0, which is the functional equivalent of RPL10 (Lan et al., 2022). In Arabidopsis and maize, UV-B light increases *RPL10* expression by about 300%, and the study of its mutants revealed that this ribosomal protein is responsible for translation during UV-B exposure (Ferreyra et al., 2010). Moreover, elongation factors of the EF-1A family have been proven to be inducible by salt stress and ABA applications in rice (Zi-Yin and Shou-Yi, 1999), and the eukaryotic initiation factor EiF4A2 can be down-regulated by dehydration in rice (Saidi and Hajibarat, 2020). This suggests that translation-related clusters can convey information about stress-specific peptide biosynthesis, in this case during UV-B exposure.

### Protein folding

Clusters 4 and 5 presented a total of 23 protein species related to protein folding (GO:0006457). Most of them were heat shock proteins (HSPs) with increased abundance, and they were likely separated into different clusters by their distinct cellular localization. While cluster 4 was predominantly associated with endoplasmic reticulum (ER) and mitochondria, cluster 5 included chloroplastic and cytosolic proteins. HSPs play a pivotal role in conferring biotic and abiotic stress tolerance in plants, functioning as molecular chaperones to defend plant cells against conformational stress (Ul Haq et al., 2019). This reinforces the evidence that high-altitude UV-B acted as a stressor, requiring the expression of numerous HSPs to maintain protein function across organelles.

### Minor clusters

Among the minor clusters, it’s noteworthy the increased abundance of three antioxidant proteins in cluster 6 (GO:0045454, “cell redox homeostasis”), namely an L-ascorbate peroxidase (APX), a thioredoxin-dependent peroxiredoxin (TPX) and a nucleoredoxin (NRX). While APXs are known key H_2_O_2_ detoxifying enzymes during abiotic stress, and particularly during UV-B exposure (Pandey et al., 2017), TPX and NRX are distinguished by possessing disulfide reduction activity. UV absorption by aromatic amino acid residues in proteins can generate free thiol groups that, when reacting together, form disulfide bonds (Neves-Petersen et al., 2012). This process may result in changes to the proteins conformation, which can affect their functions, activities, and interactions with other proteins (Banas et al., 2020). TPXs and NRXs are known to catalyze the reduction of disulfide bonds, restoring previous protein conformation and function during UV-B induced molecular stress (Marchal et al., 2014). Lastly, the reduced abundance of three tubulins in cluster 8 (GO:0007017, “microtubule-based process”), may suggest reduced cell expansion and/or proliferation (Hsiao and Huang, 2023). Even though a significant reduction in berry size or weight at harvest was not detected (Fig. S5), similar UV-B exclusion experiments installed earlier in the growing season have caused a notable reduction in berry volume (Berli et al., 2011). It is possible that the shorter span of the experimental settings was enough to down-regulate the tubulin cluster without resulting in a significant berry size reduction.

### Veraison STRING clusters

The Veraison STRING network separated into 13 main clusters with at least four nodes each, related to protein metabolism, primary metabolism, response to UV-B, secondary metabolism and intracellular transport (Fig. 5).

**Figure 5.**
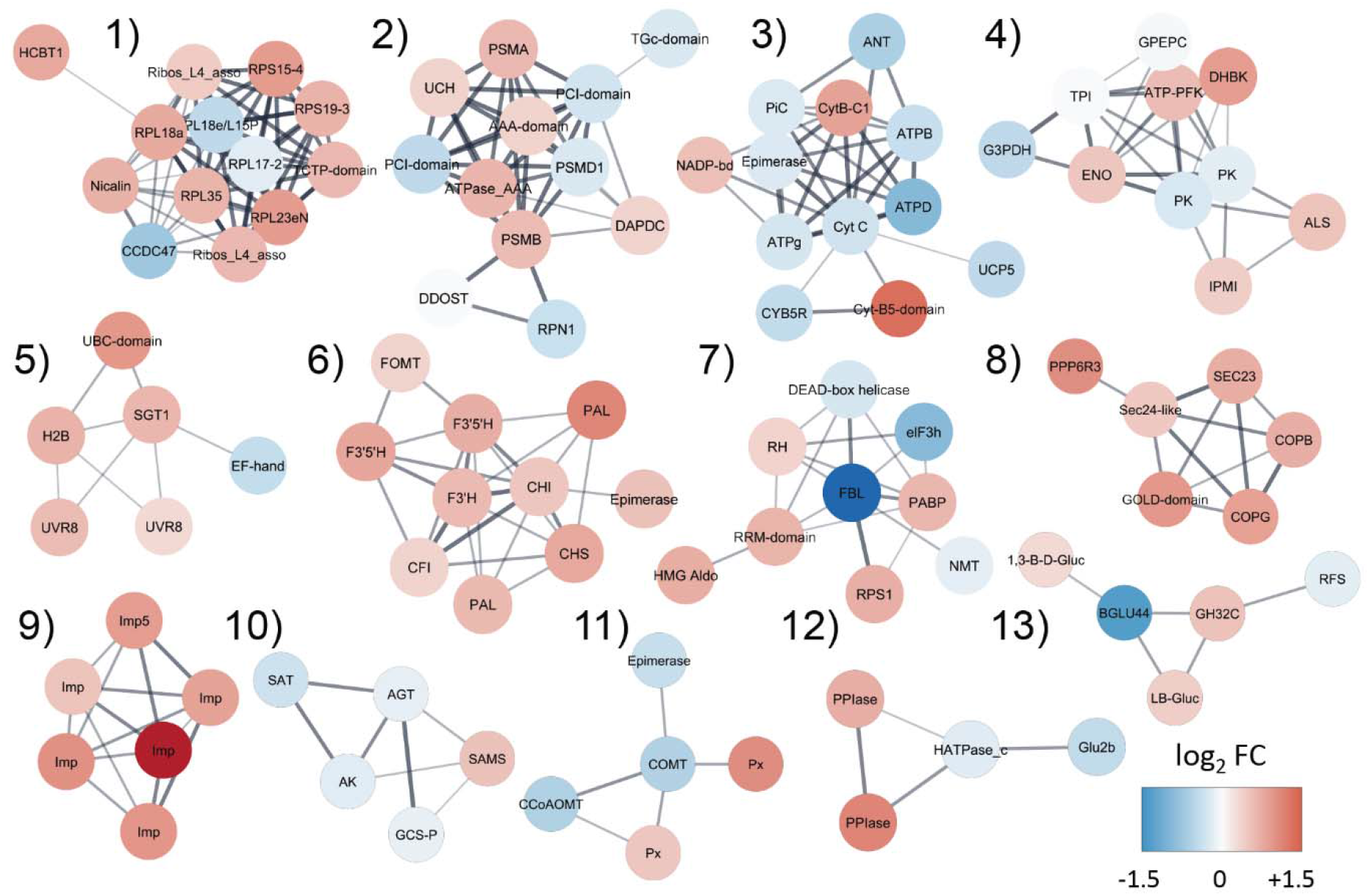
Principal STRING interaction clusters of significantly regulated proteins in veraison grape skins. The full string network was clustered using the MCL app in Cytoscape with an inflation value of 4. All clusters with at least 4 nodes are shown, colored with their log_2_ of abundance fold change as a heatmap. Representative GO terms for biological process were retrieved using STRING enrichment for each cluster with a redundancy cutoff of 0. Blue: reduced abundance; Red: increased abundance.

1. Translation
2. Protein catabolic process
3. ATP synthesis coupled proton transport
4. Glycolytic process
5. Response to UV-B
6. Flavonoid biosynthetic process
7. RNA binding
8. Endoplasmic reticulum to Golgi vesicle-mediated transport
9. Nucleocytoplasmic transport
10. Serine family amino acid metabolic process
11. Phenylpropanoid biosynthesis
12. Protein processing in endoplasmic reticulum
13. Carbohydrate metabolic process

### Peptide metabolism

Clusters 1 and 2, with a total of 25 protein species related to translation (GO:0006412) and protein catabolic process (GO:0030163), showed a marked influence of the UV-B treatment in the modulation of protein metabolism. In line with the observations made during pre-veraison, cluster 1 of veraison revealed a predominant presence of ribosomal proteins with increased abundance, likely playing a crucial role in the regulation of targeted translation processes, specifically in relationship to UV-B radiation. Cluster 2, on the other hand, was mainly composed of ubiquitin-proteasome system (UPS) structural units. The UPS is the primary proteolytic mechanism in eukaryotic cells. It consists of the initial attachment of a ubiquitin (Ub) signal to a targeted protein, and its subsequent degradation by the 26s proteasome (Stone, 2019). The proteasome comprises two parts: the 20s core unit, where degradation occurs, and the 19s regulatory unit, which itself has a base and a lid (Stone, 2019).

Interestingly, the results presented show the increased abundance of 20s core alpha and beta subunits (PSMA, D7T9I6 and PSMB, A5BYC0), and of two triple-A ATPases from the base of 19s regulatory unit (D7U9U7 and D7THJ7), but a decrease in the abundance of proteins from the lid portion of 19s unit. The less abundant 19s unit proteins were regulatory particle non-triple-A ATPases (RPNs) RPN1 (D7TM83) and PSMD1 (RPN2 homolog, D7TEW7), responsible for ubiquitin substrate recognition (Stone, 2019), and two PC1-domain proteins (F6HT17 and D7SHA5), believed to serve as scaffolds to stabilize subunits. Remarkably, the core 20s subunit can carry out Ub-independent degradation of oxidized proteins (Shringarpure et al., 2003). In fact, Kurepa et al. (2008) has demonstrated that mutating RPN proteins from the lid portion of the proteasome causes a shift from Ub-dependent to Ub-independent protein degradation in Arabidopsis, granting higher protein turnover and increased ROS tolerance. This strongly suggests that +UV-B berries modulated the architecture of the proteasome to convey a higher rate of degradation of oxidized proteins to better cope with oxidative stress.

### Primary metabolism

Cluster 3 included 12 proteins species from the mitochondrial ETC, associated with GO:0042773 “ATP synthesis coupled proton transport”. The abundances of cytochrome C (D7U0V9), and the ATP synthase subunits β, γ and δ (F6GTT2, D7SI12 and A5AY42 respectively) were reduced, as well as the inorganic phosphate carrier PiC (F6H440) and the decoupling protein UCP5 (A5C3N2), implying impaired oxidative phosphorylation. This is consistent with the ETC disruption observed in pre-veraison, including the increased abundance of the same putative alternative NAD(P)H-dehydrogenase (F6HL96).

As stated before, a reduction in the synthesis of ATP usually causes the activation of compensatory respiration pathways, such as glycolysis. And indeed, cluster 4 consists of nine proteins related to the glycolytic process (GO:0006096). The increased abundance of an enolase (A5C3N2), an ATP-dependent 6-phosphofructokinase (ATP-PFK, F6I7K1), 3,4-dihydroxy-2-butanone kinase (DHBK, D7TYA2), isopropylmalate isomerase (IPMI, F6HMP3), and acetolactate synthase (ALS, F6H3X4) suggests an increased energy demand in response to the impairment of oxidative phosphorylation caused by UV-B stress. This is supported by Kühn et al. (2015) observations of Arabidopsis mutant lines lacking complex I, in which glycolysis and TCA cycle fluxes were dramatically increased. Additionally, a significant portion of metabolites that enter glycolysis serve to fuel secondary metabolism (Plaxton, 1996), which is well known to be activated by UV-B. In regard to this catabolic demand, it is worth noting that cluster 13 includes three proteins with carbohydrate hydrolase activity that presented increased abundance: a glucan endo-1,3-beta-D-glucosidase (1,3-B-D-Gluc, E0CQB9), a beta-glucosidase (B-Gluc, F6HAB0) and a beta-fructofuranosidase (GH32C, F6HAU0). These can turn complex carbohydrates into monomers, possibly supplying glycolysis with glucose (Abbas et al., 2022; Davies and Henrissat, 1995).

### UV-B signaling

Cluster 5, labeled with the term “response to UV-B” (GO:0010224), consisted of 6 proteins related to UVR8-dependent UV-B light perception. UVR8 is the specific UV-B photoreceptor in plants (Tilbrook et al., 2013). After photoperception, UVR8 binds to chromatin through interaction with histone H2B (Cloix and Jenkins, 2008), which displayed increased abundance (F6GV40). This interaction culminates in an altered expression of a wide range of target genes, among which lies the regulation of the flavonoid biosynthesis pathway through HY5 transcription factor activation (Tilbrook et al., 2013). Intriguingly, two predicted protein species for UVR8 were detected with increased abundance: D7SWF9 and D7TT51. When comparing them with the Arabidopsis UVR8 reference sequence through BLAST alignment, the former exhibited 81.57% identity, while the latter only presented 34.29% identity (data not shown). They both did, however, display a remarkably similar AlphaFold predicted structure as compared to the X-ray crystal structure of Arabidopsis UVR8 (data not shown). It is generally established that Arabidopsis only has a single UVR8 protein, unlike *Physcomitrella patens*, which expresses two UVR8 genes, and *Marchantia polymorpha* that through alternative splicing generates two functional UVR8 species from a single gene (Soriano et al., 2018). Although it is well beyond the scope of this paper, it would be interesting to explore whether this is just a product of lacking annotation, or if Angiosperms other than Arabidopsis could have different UVR8 protein species.

Results also showed the increased abundance of an ubiquitin conjugating enzyme (UBC, D7TH48), which carries a Ub residue to be transferred to a target protein for its degradation in the proteasome (Vierstra, 2003). Interestingly, wild rice OgUBC1 is strongly induced by UV-B exposure, and its ectopic expression in Arabidopsis caused anthocyanin accumulation in leaves through the induction of chalcone synthase (*CHI*) gene. Additionally, it conferred increased tolerance to UV-B-mediated cell damage and to Botrytis sp. infection (Jeon et al., 2012). The abundance of ubiquitin ligase-associated protein SGT1 (E0CUS4) was also increased. Peart et al. (2002) showed that SGT1 is required for R protein-mediated pathogen resistance, as mutating this protein compromises the cell death involved in hypersensitive response. There is enough evidence to suggest that the UV-B response pathway shares several signaling components with the biotic stress response pathway in plants. As reviewed by Meyer et al. (2021), low doses of UV-B can boost plant defense mainly through the UVR8-mediated production of specialized metabolites, often involving the plant hormones salicylic and jasmonic acids. In their transcriptomic study, Pontin et al. (2010) showed how Malbec plantlets exposed to both low and high fluence UV-B, underwent the up-regulation of phytoalexins genes and disease resistance R genes.

### Secondary metabolism

Clusters 6 and 11 included a total of 15 proteins species that are involved in the phenylpropanoid pathway. All proteins in cluster 6 belonged to the term “flavonoid biosynthetic process” (GO:0009813) and displayed increased abundance, unequivocally indicating a complete activation of this pathway. These comprise two species of phenylalanine ammonia-lyase (PAL, A5BPT8 and F6HNF5), a chalcone synthase (CHS, Q8W3P6), two chalcone isomerases (CHI, D7T475 and F6HC36), a F3’H (A2ICC8) and two species of F3’5’H (F6HA89 and F6HA82).

The biosynthesis of flavonoids in grapes involves multiple enzymes and intermediate compounds. It starts with the conversion of phenylalanine to cinnamic acid by PAL. Cinnamic acid is then converted to p-coumaroyl-CoA by 4-coumarate:CoA ligase. The next step involves CHS forming naringenin chalcone, which is then converted to naringenin by CHI. Naringenin can be hydroxylated by F3’H to form dihydroquercetin, or by F3’5’H to form dihydromyricetin. These compounds can then be methoxylated by O-methyltransferases (OMTs) to form methoxylated derivatives (Castellarin et al., 2007). This pathway has been reported to be transcriptionally up-regulated by UV-B in Malbec plantlets (Pontin et al., 2010) and in Tempranillo grapes (Martínez-Lüscher et al., 2014), as well as in plants in general (Singh et al., 2023).

The multifunctional family of caffeoyl-coenzyme A O-methyltransferases (CCoAOMT) is ascribed to both lignin biosynthesis and stress-related anthocyanin methylation (Giordano et al., 2016). In this study, it was identified a CCoAOMT with increased abundance in cluster 6 (F6HGA0, VviAOMT2), associated by STRING with both F3’5’H isoforms, and known to harbor specific catalytic activity for the methylation of delphinidin 3-glucoside in grapevine (Fournier-Level et al., 2011). This coincides with the marked reduction in delphinidin found in +UV-B grapes (Table 2), as it is likely methoxylated into a derivative with enhanced oxidative stability. On the other hand, two CCoAOMT with reduced abundance (D7SVV2 and F6H775) were found in cluster 11 (GO:0009699, “phenylpropanoid biosynthetic process), whose orthologous are involved in lignin biosynthesis in ryegrass and petunia, respectively (Louie et al., 2010; Shaipulah et al., 2016). Moreover, the down-regulation of petunia *PhCCoAOMT1* leads to the transcriptional up-regulation of anthocyanin transcription factors, arguably due to a perturbation in the pool of secondary metabolites (Shaipulah et al., 2016).

### RNA processing

Cluster 7 included 9 proteins species involved in “RNA binding” (GO:0003723) and processing. RNA binding proteins (RBPs) interact with RNA to modulate co- and post-transcriptional events during gene regulation (Muthusamy et al., 2021). Their genes are typically up-regulated during abiotic stress, acting as RNA chaperones and directing the aggregation of mRNA transcripts into stress granules (SGs) (Yan et al., 2022). It was found that +UV-B caused the increased abundance of an RRM-domain protein (D7SKJ4), an RNA helicase (F6HPS1), a S1-domain protein (D7SYG9), a polyadenylate-binding protein (PABP, F6HQ88) and a 4-hydroxy-4-methyl-2-oxoglutarate aldolase (HMG aldolase, F6GX20). Additionally, outside of cluster 7 a minor cluster with 3 RBPs with increased abundance was detected, namely an RRM-domain protein (F6HQM5) and 2 KH-domain protein species (F6HN29 and F6H392). All of these are known to include RNA binding domains, and PABP specifically is required for SG aggregation (Yan et al., 2022).

Briefly, SGs are cytoplasmic condensates of RNA and proteins that form during stress and attune transcription to favor stress-induced transcripts in eukaryotic cells (Cabral et al., 2022). They can be categorized in canonical or non-canonical SGs by whether they respectively involve or not the eukaryotic initiation factor 3 (eIF3), which presents reduced abundance (D7U9M6) in the +UV-B condition. Moreover, non-canonical SGs aggregate during oxidative stress and UV exposure (Cabral et al., 2022). Altogether, the modulation of cluster 7 suggests that formation of non-canonical SGs could be taking place in +UV-B berries.

### Intracellular trafficking

Cluster 8, associated with the term “nucleocytoplasmic transport” (GO:0006913) was comprised of 6 importins, all with increased abundance (F6I4A5, F6HLE6, F6I6F2, A5BVQ5, D7U509 and F6GUN0). Importins mediate the nucleocytoplasmic transport of proteins by recognizing a nuclear localization signal and aiding translocation through the nuclear pore complex (Xiong et al., 2021). It is tempting to associate this cluster with the enigmatic mechanism of UVR8 internalization. However, currently there is not enough reliable data to assert that UVR8-COP1 complex translocates to the nucleus through COP1 interaction with an importin, and it rather seems like UVR8 diffuses freely through the nuclear pore (Fang et al., 2022). All these importins with increased abundance, nevertheless, could be accompanying the nuclear internalization of other transcription factors during UV-B perception or UV-related stress, as reviewed by Xiong et al. (2021).

In line with this, cluster 9, identified as “Endoplasmic reticulum to Golgi vesicle-mediated transport” (GO:0006888), displayed the increased abundance of 6 proteins species related to transport between the Golgi apparatus and the endoplasmic reticulum. These proteins are a subunit of protein phosphatase 6 (PPPP6R3, D7TKH5), two components of COPII coat complex (Sec23, F6I1E4 and Sec24, E0CT66), a GOLD-domain protein (D7UAI9) and two subunits of COPI coat complex (COPB, F6HX23 and COPG, D7TQ06). This suggests an increased need for protein trafficking and processing under higher UV-B radiation conditions, which often involves both COPI and COPII, as reviewed by Liu and Li (2019).

## Conclusions

Based on the compelling evidence presented, conclusions regarding the intricate response of Malbec grape berries to high-altitude solar UV-B radiation can be drawn. Solar UV-B can play a dual role, functioning as both a photomorphogenic signal through the UVR8 pathway and as a stressor, triggering the generation of ROS and impacting various cellular processes. Unlike artificial light sources, natural sunlight encompasses a broad spectrum of wavelengths and intensities throughout the day, allowing for a more nuanced assessment of the berries natural UV-B response. In general, exposure to +UV-B caused a strong activation of the antioxidant defense system, as is evident from the prevalence of proteins with increased abundance in the “response to hydrogen peroxide” gene ontology term and the formation of STRING clusters associated with cellular redox homeostasis. Meanwhile, proteins involved in photosynthesis and mitochondrial electron transport chain had their abundances reduced either transcriptionally or through direct UV-B. Simultaneously, proteins in charge of compensatory respiration pathways, such as glycolysis and TCA cycle, displayed increased abundance. Moreover, several cellular functions were directed towards maintaining homeostasis, with protein clusters in +UV-B indicating a widespread activation of chaperones and an increased turnover of peptides, as indicated by the regulation of ribosomal and proteasomal structures. During the veraison stage, +UV-B notably increased the abundance of proteins associated with the UVR8 signaling pathway and the phenylpropanoid biosynthetic process. This was accompanied by an overall increase in phenolic compounds, and a shift towards forms of anthocyanins with enhanced antioxidant capacity.

Ultimately, the grapevine represents a unique system in which a certain degree of stress can yield beneficial outcomes. This stress-induced response enhances biosynthesis of specialized metabolites, granting both a sounder stress resilience and a higher grape quality for winemaking.

## Notes

### Competing Interest Statement

The authors have declared no competing interest.

## Citations

Abbas, A., Shah, A. N., Shah, A. A., Nadeem, M. A., Alsaleh, A., Javed, T.,… & Abdelsalam, N. R. (2022). Genome-wide analysis of invertase gene family, and expression profiling under abiotic stress conditions in potato. Biology, 11(4), 539.

Antoniolli, A., Fontana, A. R., Piccoli, P., & Bottini, R. (2015). Characterization of polyphenols and evaluation of antioxidant capacity in grape pomace of the cv. Malbec. Food Chemistry, 178, 172–178.

Arias, L. A., Berli, F., Fontana, A., Bottini, R., & Piccoli, P. (2022). Climate change effects on grapevine physiology and biochemistry: Benefits and challenges of high altitude as an adaptation strategy. Frontiers in Plant Science, 13.

Banas, A. K., Zgłobicki, P., Kowalska, E., Bazant, A., Dziga, D., & Strzałka, W. (2020). All you need is light. Photorepair of UV-induced pyrimidine dimers. Genes (Basel), 11(11), 1304–1321.

Berli, F. J., & Bottini, R. (2013). UV-B and abscisic acid effects on grape berry maturation and quality. Journal of Berry Research. 3(1), 1–14. doi: 10.3233/JBR-130047

Berli, F. J., Alonso, R., Bressan-Smith, R., & Bottini, R. (2012). UV-B impairs growth and gas exchange in grapevines grown in high altitude. Physiologia Plantarum, 149(1), 127–140.

Berli, F. J., D’Angelo, J., Cavagnaro, B., Bottini, R., Wuilloud, R., & Silva, M. F. (2008). Phenolic composition in grape (Vitis vinifera L. cv. Malbec) ripened with different solar UV-B radiation levels by capillary zone electrophoresis. Journal of Agricultural and Food Chemistry, 56(9), 2892–2898.

Berli, F. J., Fanzone, M., Piccoli, P., & Bottini, R. (2011). Solar UV-B and ABA are involved in phenol metabolism of Vitis vinifera L. increasing biosynthesis of berry skin polyphenols. Journal of Agricultural and Food Chemistry, 59(9), 4874–4884. doi: 10.1021/jf200040z

Berli, F. J., Moreno, D., Piccoli, P., Hespanhol-Viana, L., Silva, M. F., Bressan-Smith, R.,… & Bottini, R. (2010). Abscisic acid is involved in the response of grape (Vitis vinifera L.) cv. Malbec leaf tissues to ultraviolet-B radiation by enhancing ultraviolet-absorbing compounds, antioxidant enzymes and membrane sterols. Plant, Cell & Environment, 33(1), 1–10.

Blancquaert, E. H., Oberholster, A., Ricardo-da-Silva, J. M., & Deloire, A. J. (2019). Effects of abiotic factors on phenolic compounds in the grape berry-a review. South African Journal of Enology and Viticulture, 40(1), 1–14.

Cabral, A. J., Costello, D. C., & Farny, N. G. (2022). The enigma of ultraviolet radiation stress granules: Research challenges and new perspectives. Frontiers in Molecular Biosciences, 9, 1066650.

Castellarin, S. D., Pfeiffer, A., Sivilotti, P., Degan, M., Peterlunger, E., & Di Gaspero, G. (2007). Transcriptional regulation of anthocyanin biosynthesis in ripening fruits of grapevine under seasonal water deficit. Plant, Cell & Environment, 30(11), 1381–1399.

Chen, Z., Dong, Y., & Huang, X. (2022). Plant responses to UV-B radiation: Signaling, acclimation and stress tolerance. Stress Biology, 2(1), 51.

Cloix, C., & Jenkins, G. I. (2008). Interaction of the Arabidopsis UV-B-specific signaling component UVR8 with chromatin. Molecular plant, 1(1), 118–128.

Coombe, B.G. (1995) Adoption of a system for identifying grapevine growth stages. Australian Journal of Grape and Wine Research 1, 104–110.

Davies, G., & Henrissat, B. (1995). Structures and mechanisms of glycosyl hydrolases. Structure, 3(9), 853–859.

Doncheva N.T., Morris J.H., Gorodkin J. and Jensen L.J. (2019). Cytoscape stringApp: Network analysis and visualization of proteomics data. Journal of Proteome Research, 18:623–632.

Downey, M. O., Harvey, J. S., & Robinson, S. P. (2003). Analysis of tannins in seeds and skins of Shiraz grapes throughout berry development. Australian Journal of Grape and Wine Research, 9(1), 15–27.

Fang, F., Lin, L., Zhang, Q., Lu, M., Skvortsova, M. Y., Podolec, R.,… & Yin, R. (2022). Mechanisms of UV-B light-induced photoreceptor UVR8 nuclear localization dynamics. New Phytologist, 236(5), 1824–1837.

Ferreyra, M. L. F., Pezza, A., Biarc, J., Burlingame, A. L., & Casati, P. (2010). Plant L10 ribosomal proteins have different roles during development and translation under ultraviolet-B stress. Plant physiology, 153(4), 1878–1894.

Ferreyra, S., Torres-Palazzolo, C., Bottini, R., Camargo, A., & Fontana, A. (2021). Assessment of in-vitro bioaccessibility and antioxidant capacity of phenolic compounds extracts recovered from grapevine bunch stem and cane by-products. Food Chemistry, 348, 129063.

Fournier-Level, A., Hugueney, P., Verriès, C., This, P., & Ageorges, A. (2011). Genetic mechanisms underlying the methylation level of anthocyanins in grape (Vitis viniferaL.). BMC plant biology, 11(1), 1–14.

Geisler, D. A., Päpke, C., Obata, T., Nunes-Nesi, A., Matthes, A., Schneitz, K.,… & Persson, S. (2012). Downregulation of the δ-subunit reduces mitochondrial ATP synthase levels, alters respiration, and restricts growth and gametophyte development in Arabidopsis. The Plant Cell, 24(7), 2792–2811.

Gil, M., Pontin, M., Berli, F., Bottini, R., Piccoli, P. (2012). Metabolism of terpenes in the response of grape (Vitis vinifera L.) leaf tissues to UV-B radiation. Phytochemistry, 77, 89–98.

Giordano, D., Provenzano, S., Ferrandino, A., Vitali, M., Pagliarani, C., Roman, F.,… & Schubert, A. (2016). Characterization of a multifunctional caffeoyl-CoA O-methyltransferase activated in grape berries upon drought stress. Plant Physiology and Biochemistry, 101, 23–32.

Górecka, M., Lewandowska, M., Dąbrowska-Bronk, J., Białasek, M., Barczak-Brzyżek, A., Kulasek, M.,… & Karpiński, S. (2020). Photosystem II 22kDa protein level-a prerequisite for excess light-inducible memory, cross-tolerance to UV-C and regulation of electrical signalling. Plant, Cell & Environment, 43(3), 649–661.

Hideg, É., Jansen, M. A., & Strid, Å. (2013). UV-B exposure, ROS, and stress: inseparable companions or loosely linked associates?. Trends in plant science, 18(2), 107–115.

Hsiao, A. S., & Huang, J. Y. (2023). Microtubule Regulation in Plants: From Morphological Development to Stress Adaptation. Biomolecules, 13(4), 627.

Jacoby, R. P., Millar, A. H., & Taylor, N. L. (2018). Biochemistry: stress responses and roles in stress alleviation. Annual Plant Reviews, Plant Mitochondria, 50, 227.

Jeon, E. H., Pak, J. H., Kim, M. J., Kim, H. J., Shin, S. H., Lee, J. H.,… & Chung, Y. S. (2012). Ectopic expression of ubiquitin-conjugating enzyme gene from wild rice, OgUBC1, confers resistance against UV-B radiation and Botrytis infection in Arabidopsis thaliana. Biochemical and biophysical research communications, 427(2), 309–314.

Kataria, S., Jajoo, A., & Guruprasad, K. N. (2014). Impact of increasing Ultraviolet-B (UV-B) radiation on photosynthetic processes. Journal of Photochemistry and Photobiology B: Biology, 137, 55–66.

Kolde R. pheatmap: Pretty Heatmaps [Internet] (2019) [cited 2022 Nov 18]. Available from: https://CRAN.R-project.org/package=pheatmap

Kühn, K., Obata, T., Feher, K., Bock, R., Fernie, A. R., & Meyer, E. H. (2015). Complete mitochondrial complex I deficiency induces an up-regulation of respiratory fluxes that is abolished by traces of functional complex I. Plant Physiology, 168(4), 1537–1549.

Kurepa, J., Toh-e, A., & Smalle, J. A. (2008). 26S proteasome regulatory particle mutants have increased oxidative stress tolerance. The Plant Journal, 53(1), 102–114.

Kusano, M., Tohge, T., Fukushima, A., Kobayashi, M., Hayashi, N., Otsuki, H.,… & Fernie, A. R. (2011). Metabolomics reveals comprehensive reprogramming involving two independent metabolic responses of Arabidopsis to UV-B light. The Plant Journal, 67(2), 354–369.

Lan, T., Xiong, W., Chen, X., Mo, B., & Tang, G. (2022). Plant cytoplasmic ribosomal proteins: an update on classification, nomenclature, evolution and resources. The Plant Journal, 110(1), 292–318.

Liang, T., Yang, Y., & Liu, H. (2019). Signal transduction mediated by the plant UV-B photoreceptor UVR8. New Phytologist, 221(3), 1247–1252.

Liu, L., & Li, J. (2019). Communications between the endoplasmic reticulum and other organelles during abiotic stress response in plants. Frontiers in Plant Science, 10, 749.

Louie, G. V., Bowman, M. E., Tu, Y., Mouradov, A., Spangenberg, G., & Noel, J. P. (2010). Structure-function analyses of a caffeic acid O-methyltransferase from perennial ryegrass reveal the molecular basis for substrate preference. The Plant Cell, 22(12), 4114–4127.

Marchal, C., Delorme-Hinoux, V., Bariat, L., Siala, W., Belin, C., Saez-Vasquez, J.,… & Reichheld, J. P. (2014). NTR/NRX define a new thioredoxin system in the nucleus of Arabidopsis thaliana cells. Molecular plant, 7(1), 30–44.

Marfil, C., Ibáñez, V., Alonso, R., Varela, A., Bottini, R., Masuelli, R., et al. (2019). Changes in grapevine DNA methylation and polyphenols content induced by solar ultraviolet-B radiation, water deficit and ABA spray treatments. Plant Physiology and Biochemistry, 135, 287–294. doi: 10.1016/j.plaphy.2018.12.021

Martínez-Lüscher, J., Morales, F., Sánchez-Díaz, M., Delrot, S., Aguirreolea, J., Gomès, E., & Pascual, I. (2015). Climate change conditions (elevated CO2 and temperature) and UV-B radiation affect grapevine (Vitis vinifera cv. Tempranillo) leaf carbon assimilation, altering fruit ripening rates. Plant Science, 236, 168–176.

Martínez-Lüscher, J., Sanchez-Diaz, M., Delrot, S., Aguirreolea, J., Pascual, I., & Gomes, E. (2014). Ultraviolet-B radiation and water deficit interact to alter flavonol and anthocyanin profiles in grapevine berries through transcriptomic regulation. Plant and Cell Physiology, 55(11), 1925–1936.

Martínez-Lüscher, J., Sánchez-Díaz, M., Delrot, S., Aguirreolea, J., Pascual, I., & Gomes, E. (2014). Ultraviolet-B radiation and water deficit interact to alter flavonol and anthocyanin profiles in grapevine berries through transcriptomic regulation. Plant and Cell Physiology, 55(11), 1925–1936.

Martínez-Lüscher, J., Sánchez-Díaz, M., Delrot, S., Aguirreolea, J., Pascual, I., & Gomès, E. (2016). Ultraviolet-B alleviates the uncoupling effect of elevated CO2 and increased temperature on grape berry (Vitis vinifera cv. Tempranillo) anthocyanin and sugar accumulation. Australian Journal of Grape and Wine Research, 22(1), 87–95.

Martínez-Lüscher, J., Torres, N., Hilbert, G., Richard, T., Sánchez-Díaz, M., Delrot, S., & Aguirreolea, J. (2017). Ultraviolet-B radiation and water deficit interact to alter flavonol and anthocyanin profiles in grapevine berries through transcriptomic regulation. Plant & Cell Physiology, 58(10), 1696–1712.

Meyer, P., Van de Poel, B., & De Coninck, B. (2021). UV-B light and its application potential to reduce disease and pest incidence in crops. Horticulture Research, 8.

Muthusamy, M., Kim, J. H., Kim, J. A., & Lee, S. I. (2021). Plant RNA binding proteins as critical modulators in drought, high salinity, heat, and cold stress responses: an updated overview. International Journal of Molecular Sciences, 22(13), 6731.

Negri, A. S., Prinsi, B., Rossoni M., Failla O., Scienza A., Cocucci M., (2008). Proteome changes in the skin of the grape cultivar Barbera among different stages of ripening. BMC Genomics. 8;9(1):378.

Neuhoff V., Arold N., Taube D., Ehrhardt W (1988). Improved staining of proteins in polyacrylamide gels including isoelectric focusing gels with clear background at nanogram sensitivity using Coomassie Brilliant Blue G-250 and R-250. Electrophoresis. 9(6):255–62.

Neves-Petersen, M. T., Gajula, G. P., & Petersen, S. B. (2012). UV light effects on proteins: from photochemistry to nanomedicine. Molecular photochemistry-various aspects, 125–158.

Palma, C. F. F., Castro-Alves, V., Morales, L. O., Rosenqvist, E., Ottosen, C. O., & Strid, Å. (2021). Spectral composition of light affects sensitivity to UV-B and photoinhibition in cucumber. Frontiers in Plant Science, 11, 610011.

Pandey, S., Fartyal, D., Agarwal, A., Shukla, T., James, D., Kaul, T.,… & Reddy, M. K. (2017). Abiotic stress tolerance in plants: myriad roles of ascorbate peroxidase. Frontiers in plant science, 8, 581.

Peart, J. R., Lu, R., Sadanandom, A., Malcuit, I., Moffett, P., Brice, D. C.,… & Baulcombe, D. C. (2002). Ubiquitin ligase-associated protein SGT1 is required for host and nonhost disease resistance in plants. Proceedings of the National Academy of Sciences, 99(16), 10865–10869.

Plaxton, W. C. (1996). The organization and regulation of plant glycolysis. Annual review of plant biology, 47(1), 185–214.

Pontin, M. A., Piccoli, P. N., Francisco, R., Bottini, R., Martinez-Zapater, J. M., & Lijavetzky, D. (2010). Transcriptome changes in grapevine (Vitis viniferaL.) cv. Malbec leaves induced by ultraviolet-B radiation. BMC Plant Biology, 10(1), 1–13.

R Core Team (2023). R: A language and environment for statistical computing. R Foundation for Statistical Computing, Vienna, Austria. URL https://www.R-project.org/.

Rácz, A., & Hideg, É. (2021). Narrow-band 311 nm ultraviolet-B radiation evokes different antioxidant responses from broad-band ultraviolet. Plants, 10(8), 1570.

Rai, N., Morales, L. O., & Aphalo, P. J. (2021). Perception of solar UV radiation by plants: photoreceptors and mechanisms. Plant Physiology, 186(3), 1382–1396.

Saidi, A., & Hajibarat, Z. (2020). In-silico analysis of eukaryotic translation initiation factors (eIFs) in response to environmental stresses in rice (Oryza sativa). Biologia, 75, 1731–1738.

Schertl, P., & Braun, H. P. (2014). Respiratory electron transfer pathways in plant mitochondria. Frontiers in plant science, 5, 163.

Shaipulah, N. F. M., Muhlemann, J. K., Woodworth, B. D., Van Moerkercke, A., Verdonk, J. C., Ramirez, A. A.,… & Schuurink, R. C. (2016). CCoAOMT down-regulation activates anthocyanin biosynthesis in petunia. Plant Physiology, 170(2), 717–731.

Shannon P., Markiel A., Ozier O., Baliga N.S., Wang J.T., Ramage D., Amin N., Schwikowski B., Ideker T (2003). Cytoscape: a software environment for integrated models of biomolecular interaction networks Genome Research. 13(11):2498–504

Shringarpure, R., Grune, T., Mehlhase, J. and Davies, K.J. (2003) Ubiquitin conjugation is not required for the degradation of oxidized proteins by proteasome. J. Biol. Chem. 278, 311–318.

Singh, P., Singh, A., & Choudhary, K. K. (2023). Revisiting the role of phenylpropanoids in plant defense against UV-B stress. Plant Stress, 100143.

Soriano, G., Cloix, C., Heilmann, M., Núñez-Olivera, E., Martínez-Abaigar, J., & Jenkins, G. I. (2018). Evolutionary conservation of structure and function of the UVR 8 photoreceptor from the liverwort Marchantia polymorpha and the moss *Physcomitrella patens*. New Phytologist, 217(1), 151–162.

Stone, S. L. (2019). Role of the ubiquitin proteasome system in plant response to abiotic stress. International review of cell and molecular biology, 343, 65–110.

Sweetman, C., Wong, D. C., Ford, C. M., & Drew, D. P. (2012). Transcriptome analysis at four developmental stages of grape berry (Vitis vinifera cv. Shiraz) provides insights into regulated and coordinated gene expression. BMC genomics, 13, 1–25.

Tilbrook, K., Arongaus, A. B., Binkert, M., Heijde, M., Yin, R., & Ulm, R. (2013). The UVR8 UV-B photoreceptor: perception, signaling and response. The Arabidopsis Book/American Society of Plant Biologists, 11.

Ul Haq, S., Khan, A., Ali, M., Khattak, A. M., Gai, W. X., Zhang, H. X.,… & Gong, Z. H. (2019). Heat shock proteins: dynamic biomolecules to counter plant biotic and abiotic stresses. International journal of molecular sciences, 20(21), 5321.

Urvieta, R., Buscema, F., Bottini, R., Coste, B., & Fontana, A. (2018). Phenolic and sensory profiles discriminate geographical indications for Malbec wines from different regions of Mendoza, Argentina. Food chemistry, 265, 120–127.

Van Aken, O., Giraud, E., Clifton, R., & Whelan, J. (2009). Alternative oxidase: a target and regulator of stress responses. Physiologia plantarum, 137(4), 354–361.

Van Leeuwen, C., Destrac-Irvine, A., Dubernet, M., Duchêne, E., Gowdy, M., Marguerit, E.,… & Ollat, N. (2019). An update on the impact of climate change in viticulture and potential adaptations. Agronomy, 9(9), 514.

Vierstra, R. D. (2003). The ubiquitin/26S proteasome pathway, the complex last chapter in the life of many plant proteins. Trends in plant science, 8(3), 135–142.

Wickham H., Chang W., Henry L., Pedersen T.L., Takahashi K., Wilke C., (2022). ggplot2: Create Elegant Data Visualisations Using the Grammar of Graphics [Internet]. [cited 2022 Nov 18]. Available from: https://CRAN.R-project.org/package=ggplot2

Xiong, F., Groot, E. P., Zhang, Y., & Li, S. (2021). Functions of plant importin β proteins beyond nucleocytoplasmic transport. Journal of Experimental Botany, 72(18), 6140–6149.

Yan, Y., Gan, J., Tao, Y., Okita, T. W., & Tian, L. (2022). RNA-binding proteins: the key modulator in stress granule formation and abiotic stress response. Frontiers in Plant Science, 13, 882596.

Zi-Yin, L. I., & Shou-Yi, C. H. E. N. (1999). Inducible expression of translation elongation factor 1A gene in rice seedlings in response to environmental stresses. Journal of Integrative Plant Biology, 41(8).

