## Supplementary material for "Proteomic Analysis Reveals the Molecular Pathways Responsible for Solar UV-B Acclimation in High-altitude Malbec Berries": Fig S1-S5

### Slide 1
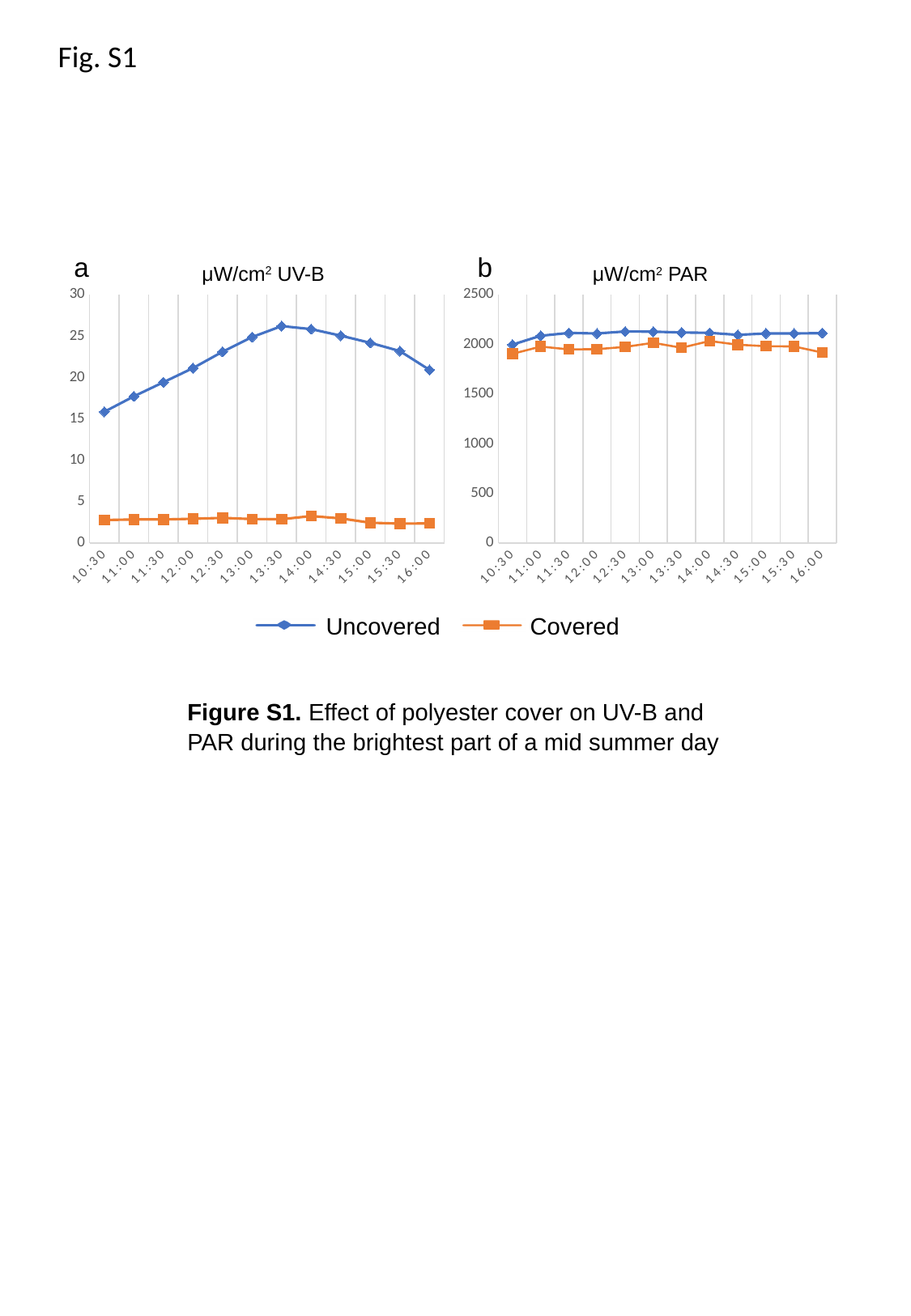

Fig. S1
a
b
μW/cm^2^ UV-B
μW/cm^2^ PAR
#### Chart
| Category | PAR + | PAR - |
|---|---|---|
| 0.4375 | 1998.0 | 1905.0 |
| 0.45833333333333331 | 2087.0 | 1978.0 |
| 0.47916666666666669 | 2115.0 | 1950.0 |
| 0.5 | 2110.0 | 1952.0 |
| 0.52083333333333337 | 2130.0 | 1975.0 |
| 0.54166666666666663 | 2128.0 | 2016.0 |
| 0.5625 | 2120.0 | 1966.0 |
| 0.58333333333333304 | 2116.0 | 2034.0 |
| 0.60416666666666696 | 2096.0 | 1997.0 |
| 0.625 | 2110.0 | 1982.0 |
| 0.64583333333333304 | 2110.0 | 1979.0 |
| 0.66666666666666596 | 2113.0 | 1918.0 |
#### Chart
| Category | UVB + | UVB - |
|---|---|---|
| 0.4375 | 15.87 | 2.78 |
| 0.45833333333333331 | 17.74 | 2.85 |
| 0.47916666666666669 | 19.44 | 2.85 |
| 0.5 | 21.15 | 2.93 |
| 0.52083333333333337 | 23.13 | 3.04 |
| 0.54166666666666663 | 24.9 | 2.9 |
| 0.5625 | 26.23 | 2.88 |
| 0.58333333333333304 | 25.86 | 3.26 |
| 0.60416666666666696 | 25.07 | 2.98 |
| 0.625 | 24.2 | 2.45 |
| 0.64583333333333304 | 23.21 | 2.35 |
| 0.66666666666666596 | 20.94 | 2.37 |Uncovered
Covered
Figure S1. Effect of polyester cover on UV-B and PAR during the brightest part of a mid summer day

### Slide 2
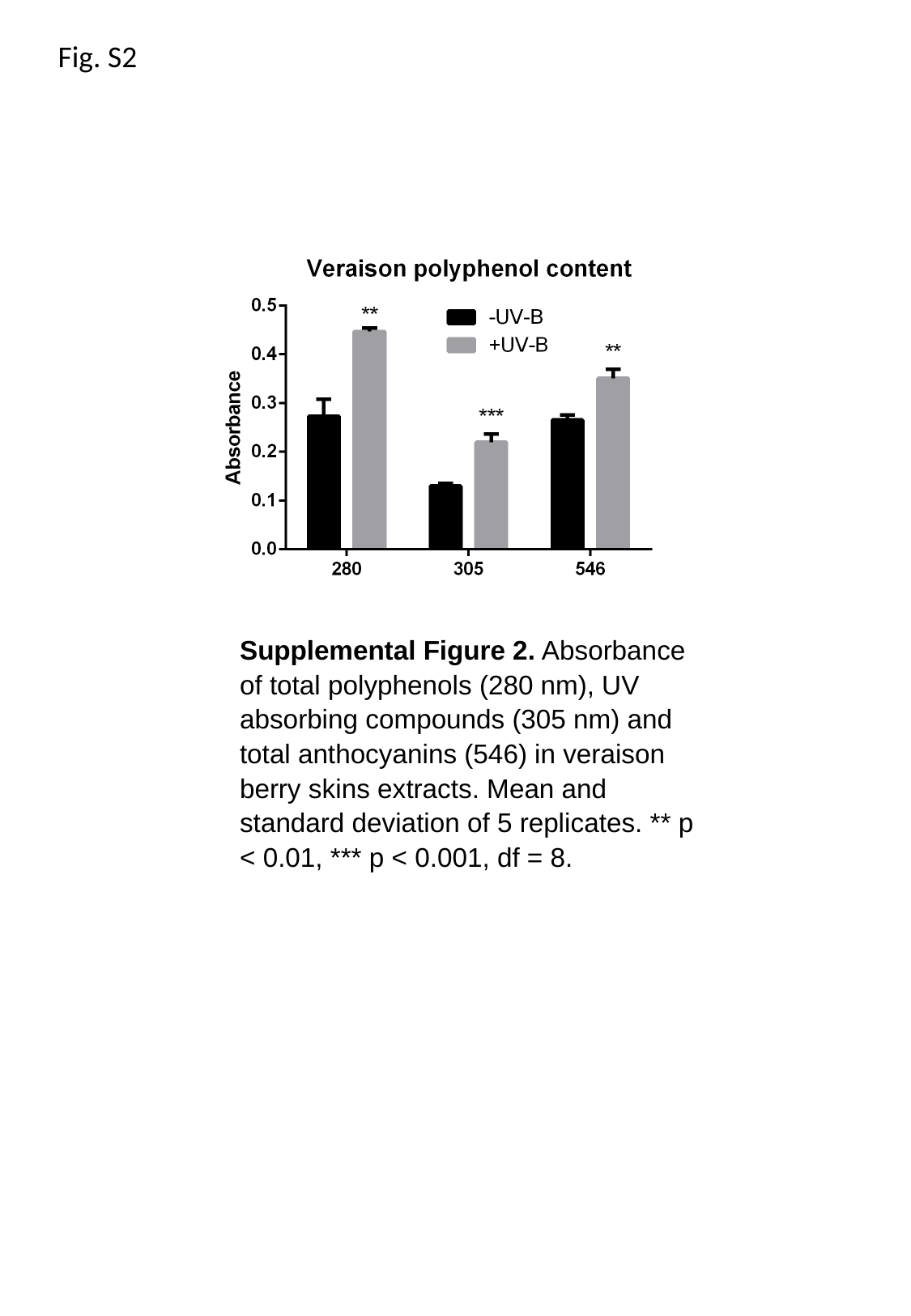

Fig. S2
Supplemental Figure 2. Absorbance of total polyphenols (280 nm), UV absorbing compounds (305 nm) and total anthocyanins (546) in veraison berry skins extracts. Mean and standard deviation of 5 replicates. ** p < 0.01, *** p < 0.001, df = 8.

### Slide 3
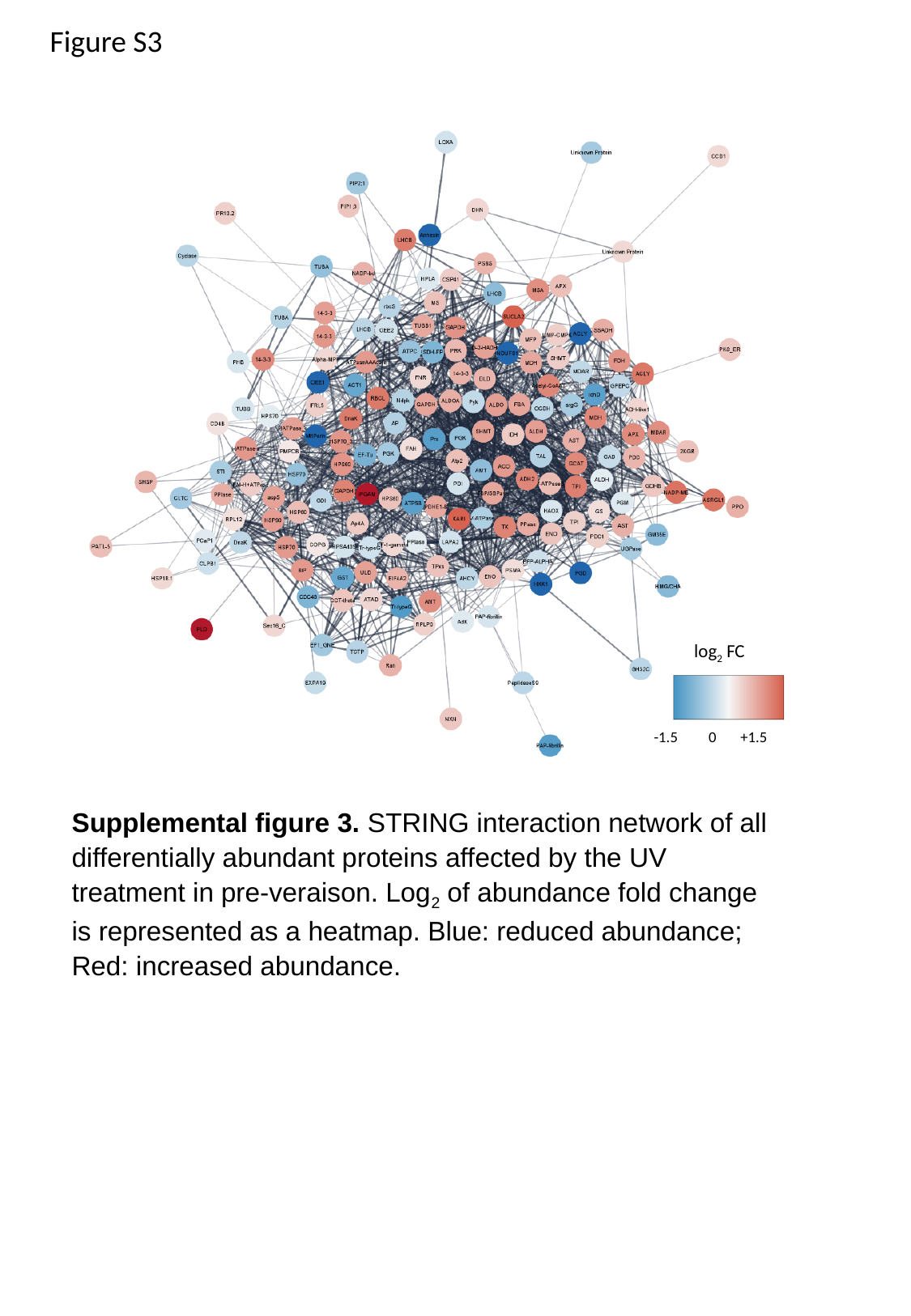

Figure S3
log_2_ FC
 -1.5 0 +1.5
Supplemental figure 3. STRING interaction network of all differentially abundant proteins affected by the UV treatment in pre-veraison. Log_2_ of abundance fold change is represented as a heatmap. Blue: reduced abundance; Red: increased abundance.

### Slide 4
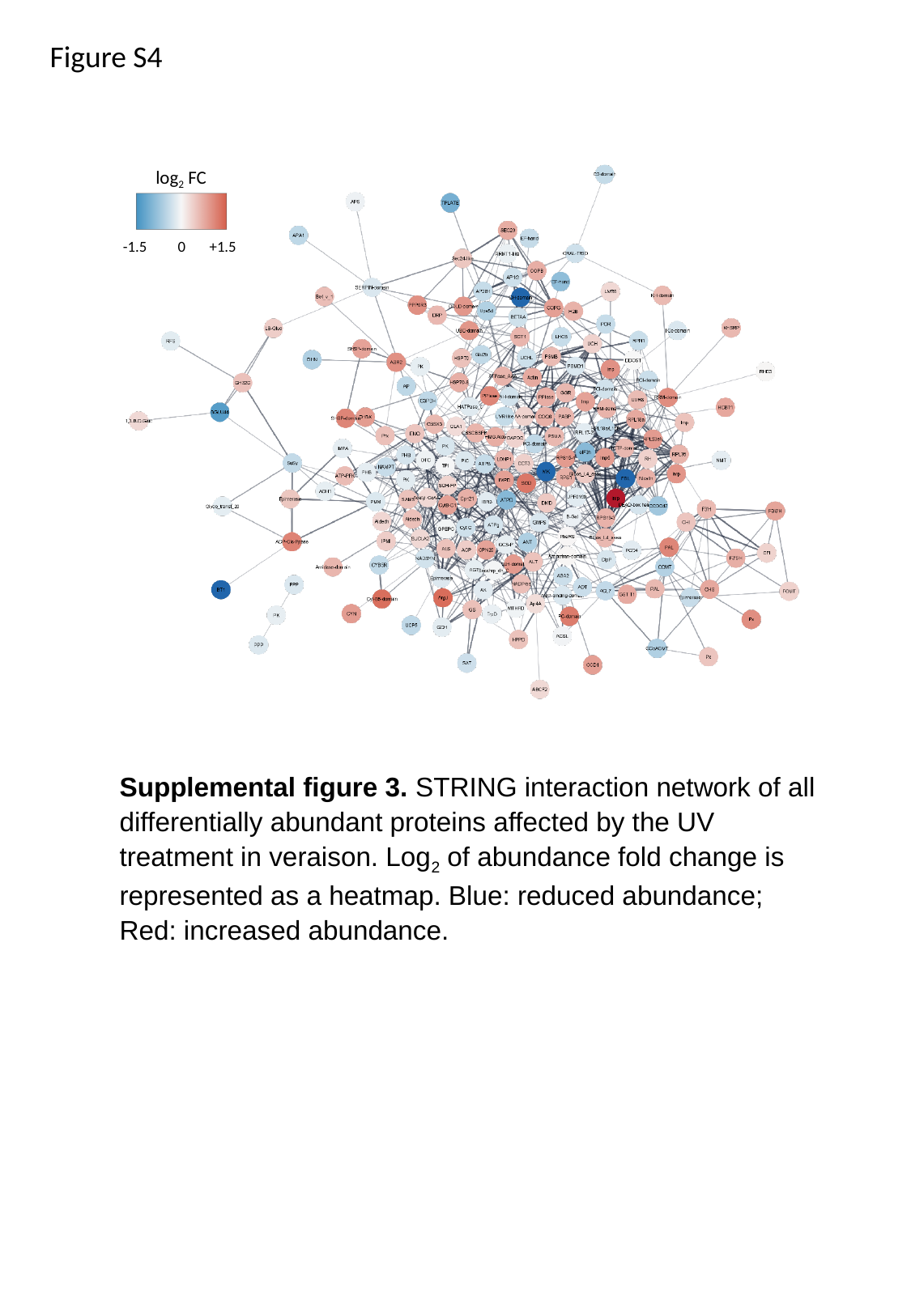

Figure S4
log_2_ FC
 -1.5 0 +1.5
Supplemental figure 3. STRING interaction network of all differentially abundant proteins affected by the UV treatment in veraison. Log_2_ of abundance fold change is represented as a heatmap. Blue: reduced abundance; Red: increased abundance.

### Slide 5
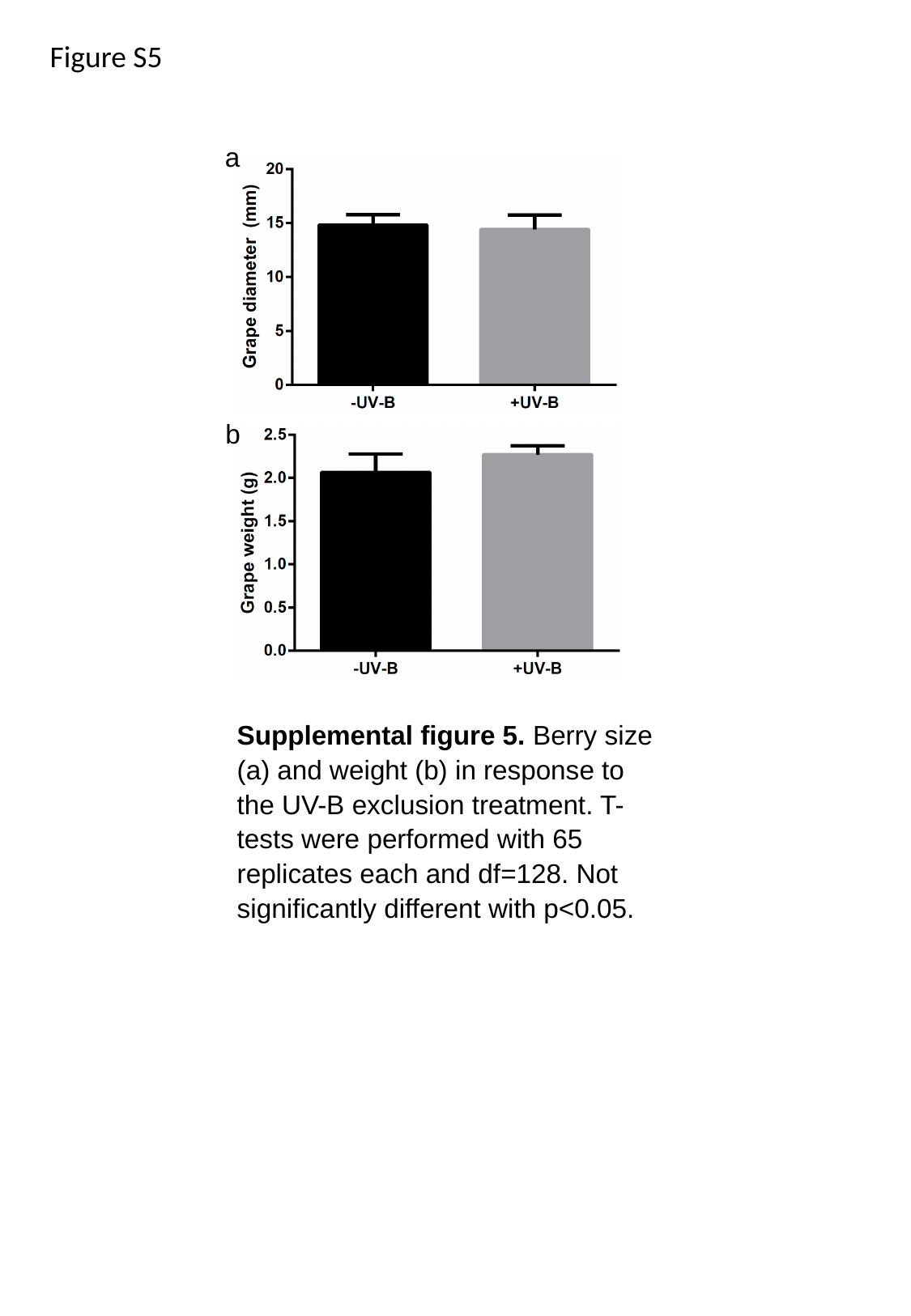

Figure S5
a
b
Supplemental figure 5. Berry size (a) and weight (b) in response to the UV-B exclusion treatment. T-tests were performed with 65 replicates each and df=128. Not significantly different with p<0.05.
